# Evaluation of Densified Collagen Tubes as an Interposition Graft in Abdominal Aorta

**DOI:** 10.64898/2026.10.07.757321

**Authors:** Alexander W. Justin, Md Abdullah Al Fahad, Federico Cammarata, Chun Ting Goh, Yifei Lai, Yiming Sun, Daniel Trajkovski, Alex Petchey, Nisha K. Ramakrishnan, Adam Winskell, Anne Segonds-Pichon, Marie-Aude Genain, Martin Goddard, Sanjay Sinha, Kourosh Saeb-Parsy, Athina E. Markaki

## Abstract

A major challenge in vascular tissue engineering is the development of scalable fabrication methods capable of producing vascular grafts with suitable mechanical properties and hemocompatibility for both preclinical evaluation and future clinical translation. Using a recently developed method to generate acellular vascular grafts from densified collagen hydrogel, we present a proof-of-concept *in vivo* evaluation of the scalable collagen graft technology. Thick-walled, genipin-crosslinked grafts, representative of a fabrication approach applicable to clinically relevant vessel sizes, were implanted as interposition grafts in the rat abdominal aorta. These grafts supported rapid neointima formation and re-endothelialisation of the luminal surface but developed late thrombosis, highlighting the challenges of evaluating clinically scalable grafts in small-animal models. To enable assessment of the platform in millimetre-scale vessels, we developed a thin-walled (∼100 µm), non-crosslinked graft using a constrained densification approach and hyaluronic acid-mediated modulation of collagen fibril assembly to improve compliance matching and hemocompatibility, with the latter providing a matrix-level approach that may complement surface-bound antithrombotic coatings. These grafts remained patent for up to 17 weeks and exhibited neointima formation, re-endothelialisation, and early cellular infiltration into the graft wall. Together, these findings demonstrate *in vivo* functionality of densified collagen vascular grafts and establish a foundation for future evaluation in larger animal models.

## 1. INTRODUCTION

Vascular grafts are required to treat a wide range of diseased and damaged blood vessels. Clinically, grafts made from synthetic materials such as expanded polytetrafluoroethylene (ePTFE) or woven polyethylene terephthalate (Dacron), which are commonly used as a replacement vessel when no biological grafting alternatives are available. However, current ePTFE and Dacron grafts suffer from high rates of occlusion – for vascular access, ePTFE grafts have a mean primary patency of 41% at 1 year[1] and cannot be remodelled by the native cells to regenerate tissue.

Bioengineered vascular grafts offer a promising alternative to currently available grafting materials, notably improving cellular response, thrombogenicity and closer compliance matching to native vessels[2,3]. One strategy involves the use of native tissue-derived grafts, in which harvested vessels are decellularised to produce ECM-based scaffolds capable of supporting recellularisation before and after implantation[4–7]. Alternatively, a wide range of biomaterials and biofabrication strategies have been employed to generate small-diameter vascular grafts *de novo;* however, despite these advances, thrombosis remains a major limitation[8]. Examples include grafts fabricated from glutaraldehyde-crosslinked, lyophilised collagen membranes[9]; carbodiimide-crosslinked, lyophilised and structured collagen/elastin[10]; electrospun, heparinised poly(L-lactide) (PLLA)[11]; polycaprolactone (PCL)[12]; cell printed matrices[13]; electrospun PCL/fibrin[14]; electrospun gelatin, PCL and genipin-crosslinked gelatin/PCL[15]; bilayered poly(L-lactide-ε-caprolactone) (PLCL)[16]; three-layered poly(lactic-co-glycolic acid) (PLGA)/polyurethane[17]; electrospun polyurethane (PU)[18]; electrospun poly(ester urea) (PEU)[19], electrospun poly(ester urethane)urea (PEUU)[20,21]; electrospun, bilayered PCL[22]; rolled sheets of collagen with elastin[23]; and air dried collagen[24].

As the main constituent component of ECM, collagen has emerged as a promising biomaterial for tissue engineering. While collagen hydrogel provides a high degree of biocompatibility, bioactivity and regenerative capability, it is mechanically weak and degrades rapidly. Collagen hydrogel densification methods increase collagen density by 20-30-fold while promoting alignment of the fibrillar network, resulting in stronger and more durable tubular material whilst still retaining regenerative potential, suitable for implantation as functional grafts (i.e. supporting blood flow). Methods to achieve collagen densification include plastic deformation[25,26], air drying[27], rotation-based water removal[28], cell compaction[29,30] and cyclic straining[31]. Recently, we reported a fully materials-based and seamless method for generating tubular grafts from densified collagen hydrogel with an ‘off-the-shelf’ capability[32]. The fabrication method is not dependent on cell seeding, bioreactor culture or decellularization, enabling rapid and inexpensive graft production, while making use of the bioactive and non-immunogenic benefits of densified collagen hydrogel.

Herein, we report a proof-of-concept evaluation of the densified collagen tubes as a small diameter vascular graft, replacing a section of the rat abdominal aorta, establishing that such grafts can be implanted and maintain patency for extended periods. We implanted thick-walled, genipin-crosslinked collagen grafts for which the method is scalable to human-scale vessels but is otherwise constrained to relatively large wall thicknesses for the vessels present in the rat. To address this, we present a variation on this densification technique enhancing its application to small diameter grafts. Using constrained densification and hyaluronic acid (HA) modulation, this produces thin-walled, non-crosslinked collagen tubes which better match the wall thickness and compliance of the rat abdominal aorta, demonstrate enhanced hemocompatibility of the collagen-based graft, and exhibit a capability for cellular repopulation into the graft walls, while still maintaining suturability and mechanical integrity for implantation as a replacement blood vessel.

## 2. RESULTS

### 2.1. Funnel densification generates thick-walled densified collagen tubes

Thick-walled densified collagen tubes at the scale of rat abdominal aortas were produced via a hybrid method. Using a previously described funnel densification method[32], which is directly scalable to human vessel dimensions, collagen gel was processed into a densified collagen tube. Briefly, water was withdrawn axially from the bottom of the gel, which was anchored by a porous membrane assembly. This pulled the collagen polymeric network downward, deforming the gel both axially and radially according to the geometry of the funnel, as shown in the timelapse images in Fig. 1a. The method requires collagen to flow between the central rod and chamber, which imposes a lower limit on attainable wall thicknesses. Consequently, additional steps were required to produce grafts suitable for use as small diameter vascular graft. To address this limitation, the densified collagen tubes were rotationally air dried following funnel densification (Fig.1b), reducing the wall thickness to ∼400 µm. Genipin crosslinking was used to stabilise the reduced wall thickness and prevent rehydration-induced swelling (Fig. 1c). Heparin was subsequently added to the graft surface to prevent clot formation. This was immobilised using a protamine sulfate treatment and crosslinked in place through an additional genipin crosslinking step[33].

**Fig 1.**
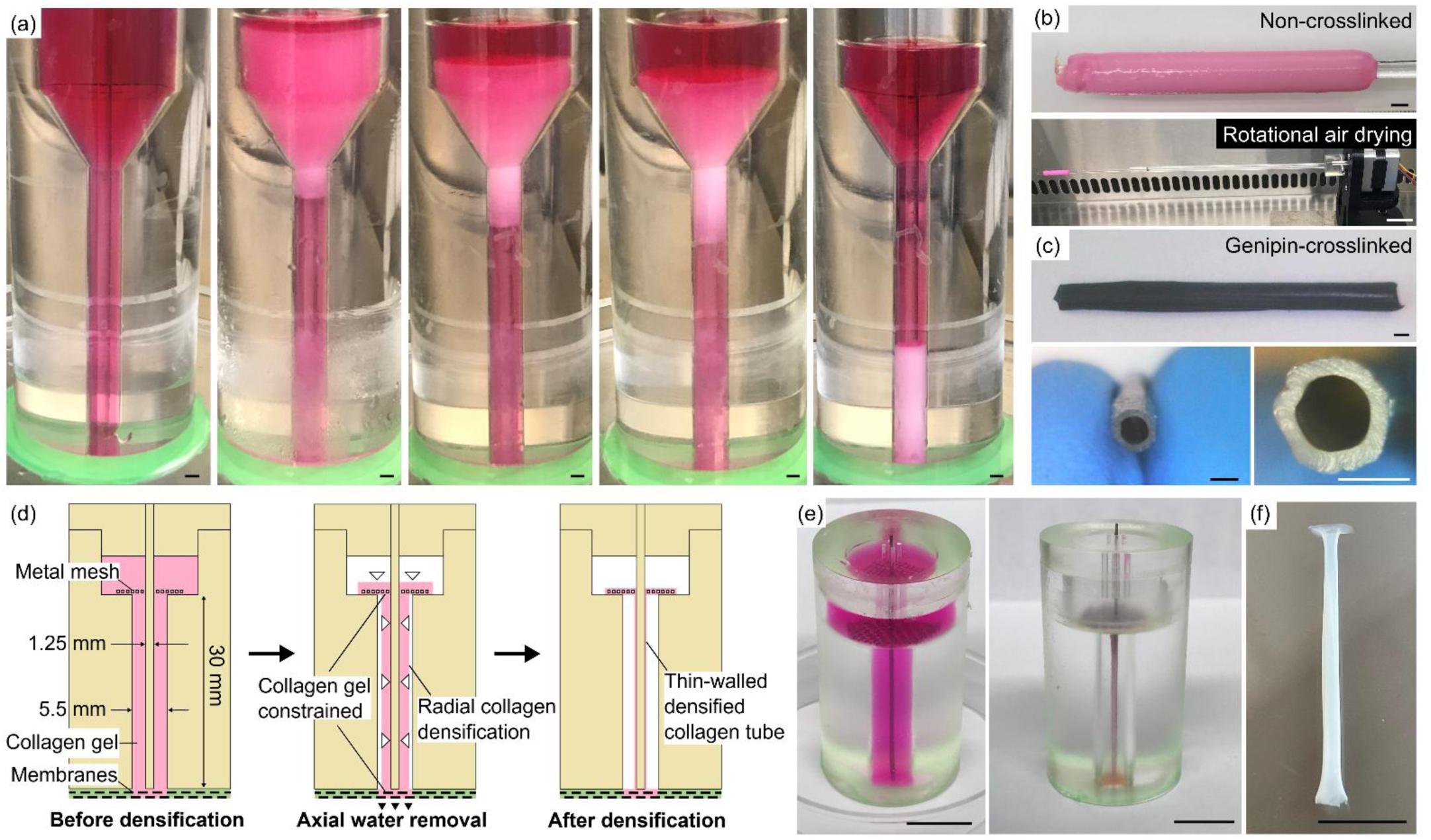
Rat-sized vascular grafts can be produced from densified collagen to be thick-walled or thin-walled.(a) Timelapse of funnel densification for thick-walled, genipin-crosslinked densified collagen tubes. **(b)** After funnel densification, tubes were air-dried to reduce wall thickness. **(c)** Genipin crosslinking of air-dried collagen tubes stabilises wall thickness and prevents rehydration-induced swelling. **(d)** Schematic of miniature funnel densification method for fabricating thin-walled, non-crosslinked densified collagen tubes. Anchored at both the top metal mesh and membranes, water removal through the base leads to radial densification of collagen hydrogel into a thin-walled collagen tube. **(e)** Images of thin-walled collagen densification method. **(f)** Resultant thin-walled densified collagen tube. Scale bars: (a, b, c): 1 mm; (e, f): 10 mm.

### 2.2. Constrained funnel densification generated thin-walled densified collagen tubes

To produce thin-walled densified collagen tubes, which more closely matched the wall thickness of the native aorta and avoided the use of chemical crosslinking needed for post-air drying stabilisation and mechanical strength, we developed a variation on the funnel densification process. In this approach, a metal mesh placed at the top of a cylindrical mould anchored the collagen gel from above, while porous membranes at the bottom anchored the gel from below (Fig. 1d). As water flows axially out of the gel through the base of the funnel, the constrained collagen polymeric network densifies radially inward (Fig. 1e, Supplementary Fig. S1a, b), producing a high density, thin-walled collagen tube (Fig. 1f). Because the collagen does not flow downwards through a small annular area between funnel and central rod, as in the larger funnel moulds, there is no constraint upon the final tube wall thickness. Therefore, wall thicknesses of ∼100 µm were achieved using this method without the need for air drying and additional stabilising crosslinking steps. Due to the non-crosslinked nature of these collagen tubes, dipyridamole (DIPY) was used as an anti-platelet coating instead of heparin.

### 2.3. Genipin crosslinking limits in vivo cell infiltration into densified collagen hydrogel

To evaluate the biocompatibility of the graft materials, disc samples of genipin-crosslinked (thick-walled) densified collagen tubes and non-crosslinked (thin-walled) densified collagen tubes were implanted subcutaneously in rats after seeding one side with smooth muscle cells (SMCs), representing a cell type expected to populate the material when placed *in situ* as a vascular graft (Fig. 2a). After 8 weeks, we found that the genipin-crosslinked collagen remained acellular (Fig. 2b). In contrast, the non-crosslinked densified collagen gels were thoroughly infiltrated by host cells (Fig. 2c). Immunostaining for the CD68 antigen identified these cells to be tissue resident macrophages, with cell densities comparable to those observed in the surrounding native tissue at 8 weeks (Fig. 2c). Staining for macrophages (CD68) and nuclei (DAPI) staining showed an increase in macrophages at later time points in relation to the graft material. This response is anticipated for a non-vascularised material and would be expected to progress toward capillary growth and matrix remodelling around the graft, thereby promoting integration with the surrounding tissue[34].

**Fig 2.**
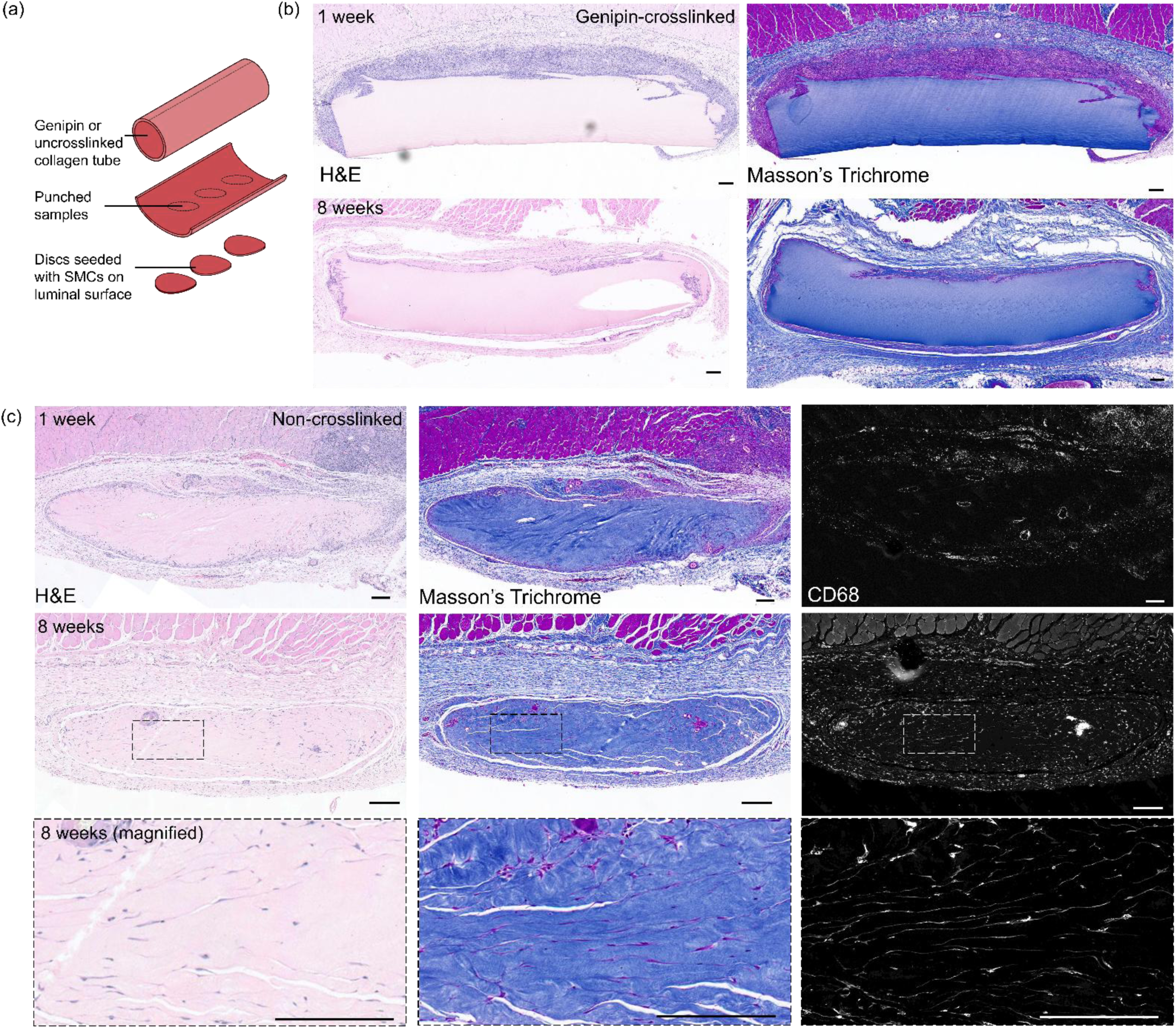
Subcutaneous implantation of densified collagen to evaluate rat in vivo biocompatibility. **(a)** Samples of densified collagen tubes, both genipin-crosslinked and uncrosslinked, were seeded with SMCs on the luminal surface to mimic in situ neointima formation. **(b)** H&E and Masson’s Trichrome staining of genipin-crosslinked densified collagen after 1 and 8 weeks showed negligible cell infiltration (*n* = 3 per timepoint). White hole in the 8-week graft is a processing artifact. **(c)** H&E, Masson’s Trichrome and CD68 immunohistochemical staining of non-crosslinked densified collagen after 1 and 8 weeks revealed significant cell infiltration (*n* = 3 per time point). CD68 staining showed infiltration of tissue resident macrophages. Collagen degradation was greater than that observed in genipin-crosslinked grafts. Scale bars: 200 µm.

### 2.4. Genipin-crosslinked, thick-walled densified collagen tubes as an interposition graft in the rat abdominal aorta

We successfully implanted genipin-crosslinked, thick-walled densified collagen tubes as an acellular vascular graft to replace a segment of the rat abdominal aorta (*n* = 8), despite a large wall thickness mismatch between the native aorta and collagen graft (Fig. 3a, b, Supplementary Video 1). After 10 weeks, the grafts remained patent and were well integrated with the anastomosed native aortic tissue (Fig. 3c, Supplementary Video 2, 3). High resolution contrast computed tomography (CT) scans performed at weeks 5-8 revealed lumen patency in all animals (Fig. 3d, Supplementary Fig. S1c). Histology (Fig. 3e) and immunostaining (Fig. 3f) revealed multi-layered neointima formation, consisting of native SMCs and a CD31+ endothelial lining. Consistent with our biocompatibility findings, no cell invasion into the walls of the genipin-crosslinked collagen was observed. However, graft extractions at 26 weeks (Fig. 3g) revealed extensive thrombus formation within the lumen of the native abdominal aorta and the vascular graft (Fig. 3h). Collagen deposition was greater in the native abdominal aorta, indicated by a darker blue staining in the lumen on MSB images. A lighter blue stain was detected in the densified collagen graft, with a clear demarcation between the native tissue and bioengineered graft. This is suggestive that thrombus formation originated in the native aorta and extended towards the anastomosis, likely due to impedance mismatch arising from compliance differences between native tissue and the genipin-crosslinked, thick-walled densified collagen graft[35–37]. Examination of the thrombus in both longitudinal and cross-sections revealed the porous nature of the clot (Fig. 3i), suggesting ongoing remodelling of an older thrombus. Mature thrombi are known to undergo cellular infiltration and fibrinolysis with subsequent extracellular matrix deposition and reorganisation, leading to clot permeability[38–40].

**Fig 3.**
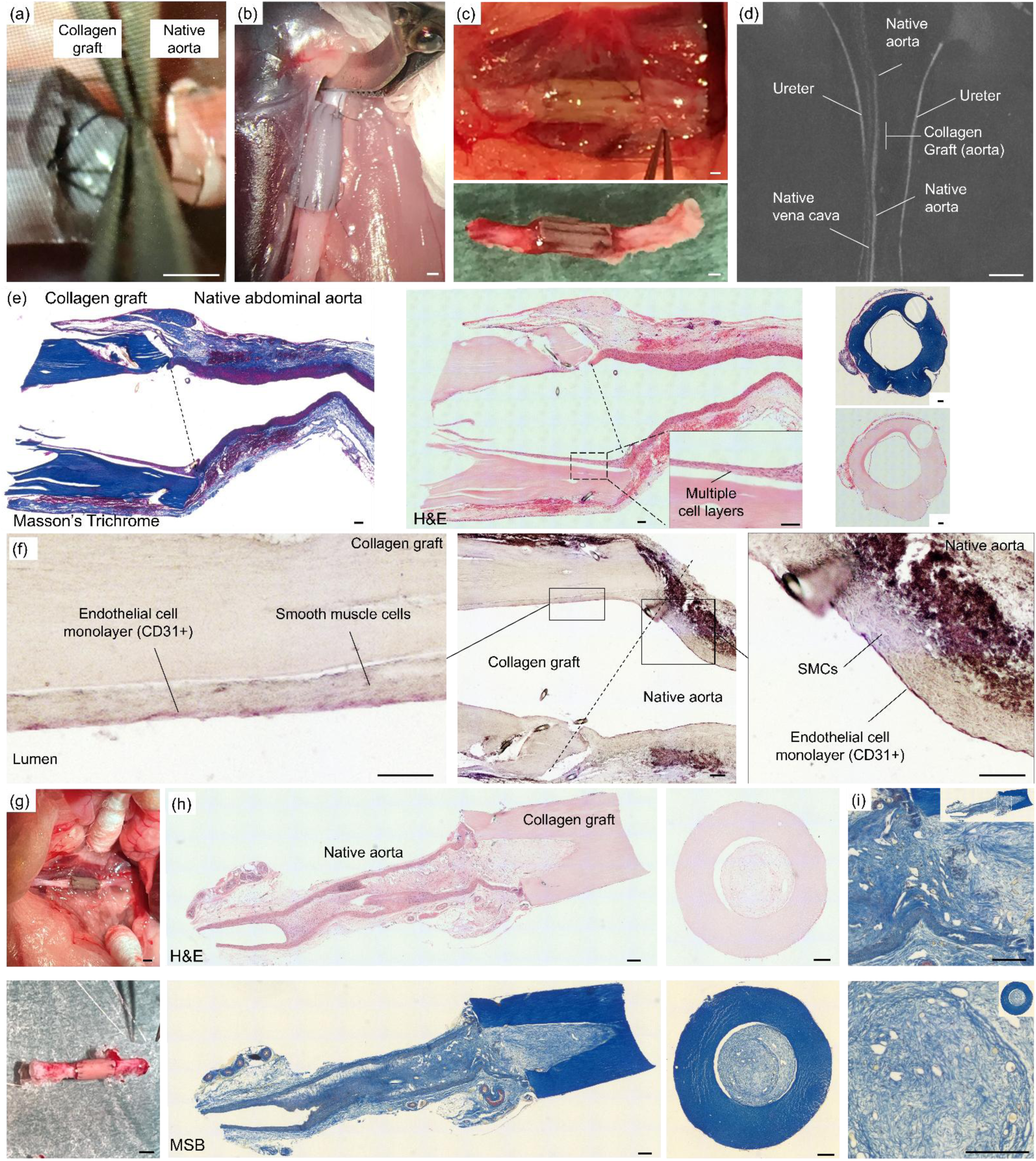
Abdominal aorta replacement in a rat model using genipin crosslinked, thick-walled collagen graft. **(a)** In situ abdominal aorta replacement showing wall thickness mismatch between graft and native abdominal aorta. Scale bar: 500 µm. **(b)** Genipin-crosslinked densified collagen tube anastomosed end-to-end with the rat abdominal aorta. Scale bar: 500 µm. **(c)** In situ collagen graft before and after extraction, with peri-anastomotic native aorta. Scale bars: 500 µm. **(d)** *In vivo* dorsal plane post-contrast maximum intensity projection (MIP) CT image of an interposition collagen graft in the rat abdominal aorta at 8 weeks post-implantation. The contrast column is intact, demonstrating patency (Rat 4). Scale bar: 5 mm. **(e)** Masson’s trichrome and H&E staining of longitudinal sections of the abdominal aorta and collagen graft anastomosis at 10 weeks. Inset shows multilayered neointima formation on the graft luminal surface. Scale bars: 100 µm. **(f)** CD31 immunohistochemical staining of longitudinal sections of the anastomosis between abdominal aorta and collagen graft showing neointima formation and re-endothelialisation of the graft luminal surface at 10 weeks. Scale bars: 100 µm. **(g)** Anastomosed collagen graft at 26 weeks in situ and after explantation. Scale bars: 1 mm. **(h)** H&E and Martius Scarlett Blue (MSB) staining of sections from the anastomosis between abdominal aorta and collagen graft, showing clot formation at 26 weeks. In the MSB stain, darker blue staining is evident in the native abdominal aorta relative to the collagen graft. Scale bars: 200 µm. **(i)** High magnification image of thrombus structure in longitudinal (top) and cross-section (bottom). As shown in (h), the longitudinal section reveals a sharp transition of dark and light blue staining in the original lumen, suggesting clot formation at the interface between graft and native vessel. Scale bars: 200 µm.

### 2.5. Thin-walled and DIPY-treated collagen tubes exhibit suitable mechanical compliance and hemocompatibility

The constrained funnel densification method enables the production of very thin-walled tubes, resulting in very high final collagen densities suitable for suturing without the need for genipin crosslinking. The wall thickness was further reduced through the addition of HA to the collagen gel precursor solution (Fig. 4a), from 138 ± 15 to 96 ± 15 µm, with a proportional increase in the collagen density, from 185 ± 21 to 278 ± 49 mg/mL.

**Fig 4.**
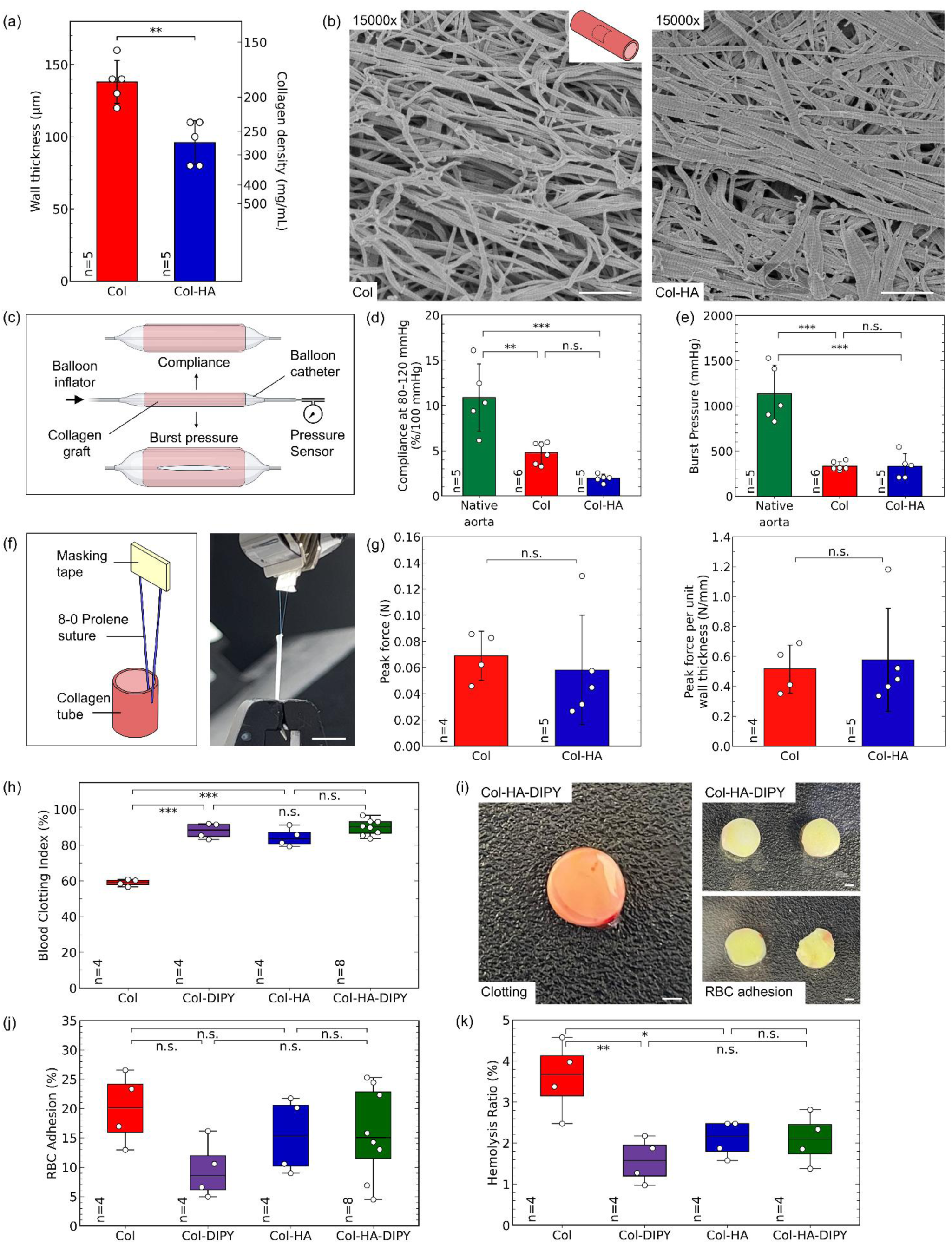
Mechanical and thrombogenic analysis of uncrosslinked, thin-walled collagen tubes with DIPY coating. **(a)** Comparison of wall thickness and corresponding collagen density of collagen and collagen with HA grafts. Col: collagen, Col-HA: collagen modulated with HA. **(b)** SEM micrographs of the outer surface of the densified collagen grafts. Scale bars: 1 µm. **(c)** Schematic of compliance and burst pressure testing configuration. **(d)** Compliance measurements of native rat abdominal aorta, collagen graft, and collagen with HA graft. **(e)** Gauge burst pressure comparison of collagen and collagen with HA grafts to native rat abdominal aorta. **(f)** Suture retention experimental setup. **(g)** Suture retention comparing collagen and collagen with HA grafts; peak force (left) and peak force per unit wall thickness (right). Scale bar: 1 mm. **(h)** Comparison of thrombogenicity of collagen and collagen with HA grafts with and without dipyridamole (DIPY) coating. Images of **(i)** clotting behaviour and **(j)** RBC adhesion on collagen-HA grafts. **(k)** Comparison of RBC adhesion of collagen and collagen with HA grafts with and without dipyridamole coating. Scale bars: 1 mm. **(l)** RBC hemolysis ratio of collagen and collagen with HA grafts with and without dipyridamole coating. Statistics: ANOVA with Tukey’s honestly significant difference (HSD) post-hoc comparisons. *p* < 0.05 (*), *p* < 0.01 (**) and *p* < 0.001 (***); n.s. means no statistically significant difference between groups was detected.

HA-containing collagen gels exhibited greater collagen fibril bundling, consistent with previous work (e.g. [41,42]). This produced a fibrillar architecture characterised by densely packed fibril bundles separated by larger interfibrillar void spaces. Such an architecture may facilitate greater volumetric reduction during densification, as collagen bundles can be rearranged and compacted more effectively than networks composed of thinner fibril bundles. Consequently, the hydrogel can be densified into a smaller final volume, producing thinner-walled grafts with higher collagen concentrations. In contrast, unmodulated collagen hydrogels formed thinner fibril bundles with smaller interfibrillar void spaces, which may limit the degree of volumetric compaction achievable during densification and result in thicker-walled grafts. Differences in fibril bundling and void morphology were observed using scanning electron microscopy (SEM) of the final densified collagen grafts (Fig. 4b, Supplementary Fig. S2), indicating that mesoscale collagen architecture influences the densification behaviour of the hydrogel.

The compliances of collagen tubes were compared to native rat aorta, with matched luminal diameters (Fig. 4c, d). Collagen tubes with and without HA exhibited compliances of 1.96 ± 0.44 and 4.80 ± 1.19 %/100 mmHg, respectively, although we did not detect a statistically significant difference. Native rat aortae of the same diameter yielded a compliance of 10.88 ± 3.70 %/100 mmHg. Although fibrils are predominantly aligned in the axial direction in collagen tubes, HA-induced fibril bundling and increased densification may increase resistance to small strain circumferential deformation under pressurisation. This may offset the increase in compliance that would otherwise be expected from the reduced wall thickness of HA-collagen tubes. We also measured the compliance of clinically available ePTFE grafts as 0.52 ± 0.15 %/100 mmHg (*n* = 8). However, the available ePTFE grafts had a substantially larger diameter (6 mm) and required a different testing methodology, limiting direct comparison with the experimental grafts.

The implanted grafts exhibited burst pressures sufficient to withstand physiologic pressures in the rat aorta, with no significant differences observed between collagen tubes with and without HA (333 ± 138 mmHg (gauge) and 335 ± 47 mmHg (gauge), respectively, Fig.4e). Native aorta, however, displayed significantly higher burst pressures (1136 ± 315 mmHg (gauge)). Although the HA-modulated collagen grafts had a higher collagen concentration from greater volumetric compaction, these changes did not translate into increased burst pressure over the non-modulated variant. Collagen fibril bundling may create regions of stress localisation, lowering failure resistance between fibril bundles. Consequently, the ultimate strength governing burst failure was not altered.

The collagen grafts displayed sufficient suture pullout forces (Fig. 4f) for implantation, with or without HA (0.058 ± 0.042 N and 0.069 ± 0.019 N, respectively). The corresponding thickness-normalised values were 0.58 ± 0.34 and 0.52 ± 0.16 N/mm, respectively (Fig. 4g). Neither the absolute suture pull-out force nor the thickness-normalised values differed significantly between the two groups. The higher variability observed in the HA-modulated group was driven by one biological replicate, with both corresponding technical replicates showing similar responses. The higher collagen density and bundled fibrillar architecture of the HA-collagen tubes may nevertheless have compensated for their reduced thickness, resulting in comparable performance between the two groups.

We also tested the hemocompatibility of the thin-walled, uncrosslinked grafts. Interestingly, the inclusion of HA into the collagen graft resulted in a blood clotting resistance significantly higher than collagen alone and comparable to a DIPY-coated collagen graft (Col: 59 ± 2 %, Col-HA: 84 ± 5 %, Col-DIPY: 88 ± 4 %). This may enable a scaffold-intrinsic protection to supplement the anti-platelet DIPY coating when used in combination (Col-HA-DIPY: 87 ± 4 %, Fig. 4h, i – left). HA exhibits high hydrophilicity which may reduce interactions between blood components and the collagenous matrix, which is well established as a pro-hemostatic substrate, through platelet adhesion and activation[43–45]. No significant differences were detected in red blood cell (RBC) adhesion between Col-DIPY, Col-HA and Col-HA-DIPY groups (Fig.4h, i – right, j), but HA addition provided a scaffold-intrinsic protection against RBC hemolysis ratio (Col: 3.6 ± 0.9 %, Col-HA: 2.1 ± 0.5 %), as shown in Fig. 4k.

Non-modulated collagen grafts exhibited a higher mean compliance than HA-modulated grafts (Δ = 2.8 %/100 mmHg), although we did not detect a statistically significant difference. Nevertheless, Collagen-HA grafts were selected for implantation owing to their intrinsic hemocompatibility and the avoidance of potential risks associated with long-term durability of the DIPY coating during blood exposure.

### 2.6. Non-crosslinked, thin-walled densified collagen tubes as interposition grafts in the rat abdominal aorta

The thin-walled densified collagen grafts (with HA) were successfully implanted acellular (*n* = 5) as a replacement for a segment of the rat abdominal aorta (Fig. 5a). These grafts more closely matched the wall thickness of the native abdominal aorta, and their greater ability to expand, relative to the thick-walled, genipin crosslinked grafts, was visibly apparent upon re-establishment of flow. At 17 weeks post-implantation, the vascular grafts had integrated well into the native aortic tissue (Fig. 5b, Supplementary Video 4). Perfusion testing using a syringe and needle confirmed lumen patency was maintained (Fig. 5c, Supplementary Video 5). Longitudinal sectioning and H&E staining revealed multi-layered neointima formation within the lumen of the graft, together with adventitial tissue formation on the external surface of the densified collagen graft (Fig. 5d, Supplementary Fig. S3a, b). Immunostaining for αSMC and CD31 revealed the structure of the neointima, comprising multi-layered SMCs and an endothelial monolayer (Fig. 5e). Furthermore, CD31+ cells were identified within the densified collagen walls of the graft. Neointima thickness was ∼40 µm along the full length of the lumen surface (Fig. 5f). CD68, CD163, and CD45 immunostaining showed scattered instances of infiltration into the walls of the collagen graft (Fig. 5g, Supplementary Fig. S3c), aligning with the findings from the biocompatibility study.

**Fig 5.**
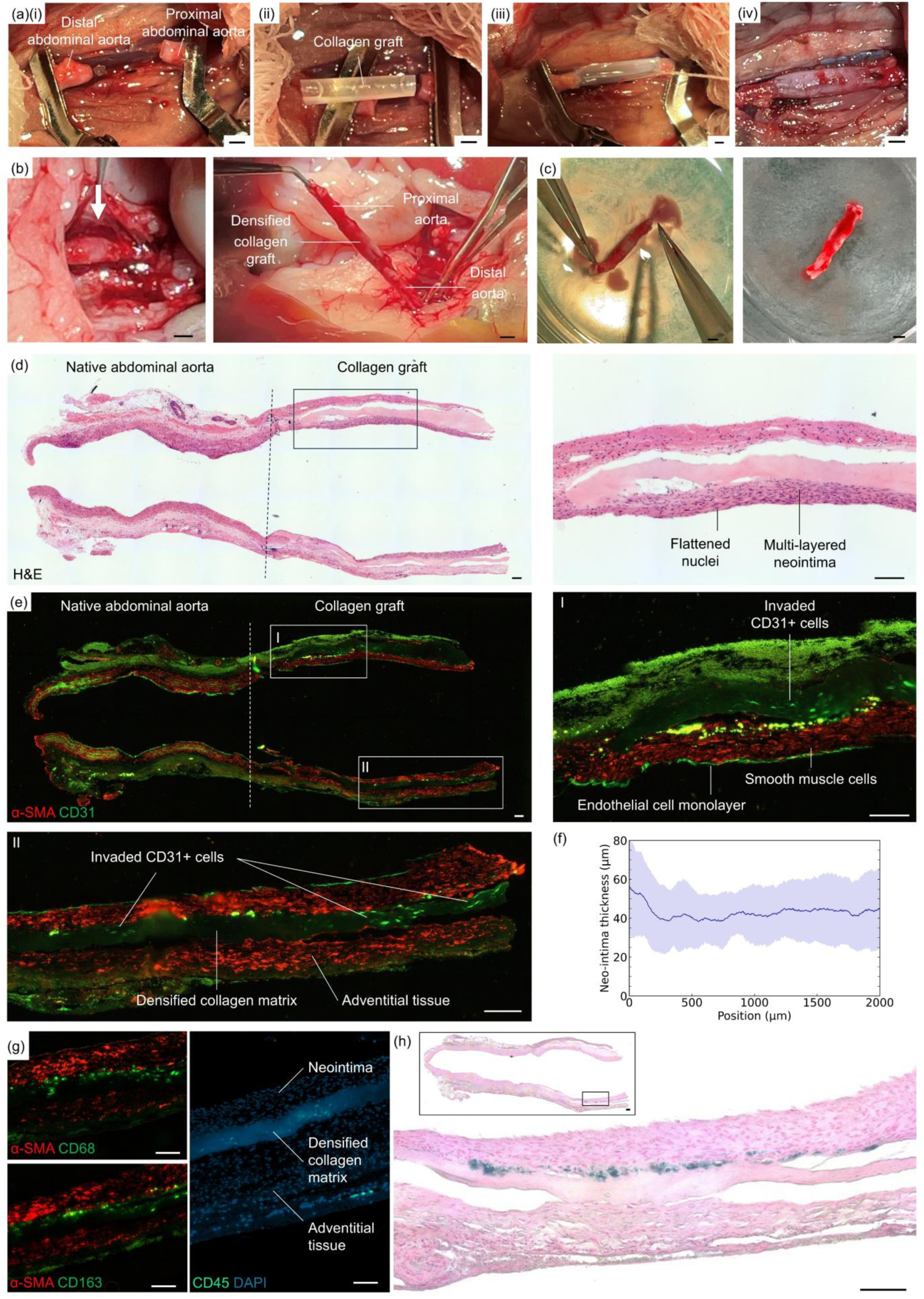
Abdominal aorta replacement in a rat model using non-crosslinked, thin-walled collagen graft. **(a)** Graft implantation. **(i)** Segment of the native abdominal aorta isolated with vascular clamps. **(ii)** Collagen graft placed adjacent to the native vessel, demonstrating a close match in lumen diameter and wall thickness. **(iii)** End-to-end anastomosis completed with sutures, with clamps still in place. **(iv)** Final view post-implantation following clamp removal, showing a stable and patent graft with no visible leakage. Scale bars: 1 mm. **(b)** Graft in situ after 17 weeks. Scale bars: 1 mm. **(c)** Upon extraction, the graft was perfused to demonstrate lumen patency. Scale bars: 1 mm. **(d)** H&E staining of a longitudinal section (left, whole section; right, magnified region) of the abdominal aorta and collagen graft showing multi-cell layered neointima formation at 17 weeks, along with flattened nuclei characteristic of an endothelium. Scale bars: 100 µm. **(e)** Immunohistochemical staining of a longitudinal section of the abdominal aorta and collagen graft showing that the neointima consists of smooth muscle layers and endothelium at 17 weeks. Adventitial tissue was present on the outer graft surface with infiltrating CD31+ cells visible in magnified regions. Scale bars: 100 µm. **(f)** Neo-intima thickness from anastomosis (0 µm) to graft mid-point (mean ± s.d., *n* = 14 sections). **(g)** CD68, CD163 and CD45 immunostaining shows sporadic infiltration of macrophages into walls of the graft. **(h)** Perls histological staining revealed limited iron deposits from RBC hemolysis prior to neointima formation. Scale bars: 100 µm.

Grafts showed a neo-adventitia around the graft which forms as part of the normal reparative process (i.e. not pathological). We observed an external, collagen-rich fibrous neo-adventitia, containing fibroblasts and capillaries (RBC-filled lumen with flattened endothelial cells), as is typical of small diameter vascular grafts[7,46]. A cell-rich neointima had formed in grafts, consisting of flattened endothelial cells (morphology and CD31-positive) and myofibroblast-like cells underneath (morphology and αSMA-positive). Cells inside the graft display a different morphology, texture, and structure to those outside. Blood flow pulsatility supports smooth muscle differentiation of the migrated myofibroblasts to a more mature contractile phenotype[47,48].

Perls staining identified limited iron deposits between the collagen graft and neointima, indicating RBC hemolysis associated with surface blood clotting prior to luminal recellularization (Fig. 5h). Although the dipyridamole coating appears to be effective in reducing platelet adhesion[49,50], complete prevention of surface clotting is unlikely, and these findings are therefore not unexpected. More modern anticoagulant drugs used clinically, such as Rivaroxaban or those of the same class, may further inhibit this surface clotting, if necessary[51]. Rivaroxaban is a direct inhibitor of activated Factor X though we do not currently know whether it can be bound to a collagen graft surface.

Overall, grafts demonstrated similar neointima formation, re-endothelialisation and healing; however, one graft showed an old, remodelled clot structure, fully occluding the lumen. Our analysis was that a technical surgical failure or graft detachment had taken place likely within the first few weeks, owing to a clear break in tissue in the graft longitudinal section and giant cell reaction visible around suture sites. New luminal structures were visible in the bulk of the clot, as is typically observed in pathological conditions.

## 3. DISCUSSION

A useful feature of a method for producing vascular grafts is its scalability across a wide range of diameters and wall thicknesses, enabling its application to different tubular tissues and evaluation in both small and large animal settings, prior to clinical application. The funnel densification technique used here to fabricate thick-walled densified collagen tubes can generate vascular grafts with luminal diameters comparable to rat abdominal aorta (1 mm) and we have previously shown[32] the same technique is applicable to fabricating vascular grafts with lumen diameters of 5 mm. However, producing vascular grafts with wall thicknesses comparable to rat blood vessels (i.e. ∼100 µm) is not feasible with this technique, even with the accompanying air-drying step. During densification, collagen and water must flow through the annular space between rod and funnel; however, aligning the central rod to within <100 µm tolerances is impractical, and frictional forces prevent collagen flow through such narrow gaps. For small diameter vascular grafts, the thin-walled densified collagen tube method detailed here is more applicable, but such a technique is not readily scalable to large animal and human-scale vascular grafts. The constrained funnel densification method does, however, allow for ultra-fine lumen diameters and wall thicknesses, which may be adapted for other tubular tissues. For example, when fabricating nerve guides that require very fine lumen diameters (10-100 µm), the central rod may be replaced with a metal wire, which is centred by passing it through a hole in the inner membrane and pulling it taut before attachment of the outer membrane. Appropriate modification of the chamber diameter and starting collagen concentration should yield the desired wall thickness for such tubes. A further reduction in wall thickness and increase in collagen density was achieved by incorporating HA which enhanced collagen fibril bundling and increased volumetric compaction during densification without compromising burst pressure or suture retention. These findings suggest that the modulation of collagen fibrillogenesis, when combined with collagen densification, provides a versatile route for engineering dense collagen constructs.

Collagen is widely recognised as a highly-thrombogenic substrate that promotes platelet adhesion, activation, and thrombus formation[52]. To counter this, we applied both an anti-platelet coating of DIPY to the surface of the graft, and an intrinsic strategy to modulate collagen fibril assembly using HA[41]. Notably, HA modulation alone substantially improved resistance to thrombus formation, approaching the effect achieved with DIPY coating. The fibrillar structure of collagen has previously been shown to strongly influence platelet adhesion and thrombus formation[53], including observations that structurally abnormal collagen fibrils in abdominal aortic aneurysm tissue exhibit reduced platelet adhesion[54]. To our knowledge, this is the first report demonstrating that deliberate modulation of collagen fibril assembly can be leveraged to improve the hemocompatibility of a collagen-based vascular graft.

Neointima formation and re-endothelialisation of the lumen at 10 weeks was very encouraging, in which SMCs and endothelial cell types had grown along luminal surface from native abdominal aorta. Comparing thick-walled, genipin-crosslinked grafts with thin-walled, non-crosslinked grafts highlights the importance of mechanical compatibility with the host vessel. Although both graft types demonstrated early patency and supported neointima formation, late thrombus formation was observed in the thick-walled, genipin-crosslinked grafts. This is likely attributable to the greater compliance mismatch between the native vessel and the thick-walled, genipin-crosslinked collagen graft. This abrupt transition can alter pulsatile hemodynamics by creating an impedance mismatch at the graft-artery interface, promoting pressure wave reflections and local disturbances in pressure, flow, and shear stress that may disrupt the endothelial cell lining at the anastomosis and provide sites for thrombus formation. Ischemic injury may also be a factor when suturing blood vessels as the endothelial cell lining in the vessel is deprived of oxygen and undergoes inflammation under re-perfusion[55]; thus, deterioration of the cell lining via necrosis[56] exposes underlying ECM and can also lead to clot formation. This may be a scale-dependent effect which may or may not occur in large animal or human-scale vascular grafts.

As is common across small diameter grafts, implantation at this scale presents significant technical challenges that constrain both reproducibility and translational relevance to larger animal models. The present study was designed to establish proof-of-concept that densified collagen tubes can be manufactured and function as vascular grafts *in vivo,* as evidenced by maintained patency and recellularisation over the study period. Collectively, these findings provide sufficient evidence of feasibility, and further small-animal experimentation is unlikely to yield additional mechanistic or translational insight. Accordingly, these results provide a robust foundation for progression to large animal studies.

## 4. CONCLUSIONS

In this study, we demonstrated the feasibility of implanting acellular densified collagen grafts generated using funnel densification methods and implanted into small animal models. Thick-walled, genipin crosslinked grafts supported neointima formation and luminal re-endothelialisation, but were found to induce clotting at later timepoints, likely owing to compliance mismatch and scale-specific limitations associated with small diameter grafts. In contrast, a new variation on the method which produces thin-walled densified collagen grafts was also tested, which exhibited improved compliance matching to the native abdominal aorta. Incorporation of HA further enhanced hemocompatibility and these grafts remained patent to 17-week timepoints. These findings highlight the importance of mechanical compatibility in performance of vascular grafts and establish densified collagen methods as a promising strategy for small diameter vascular substitutes.

## METHODS

### Fabrication of genipin-crosslinked, thick-walled densified collagen tubes

The funnel densification method was employed as previously described[32]. A 1 mm hole was made in a 1.2 µm membrane using a biopsy punch. The polycarbonate funnel was oiled and a pair of porous (1.2 µm) membranes fixed to the bottom using silicone sealant, with the holed membrane placed closest to funnel. A collagen gel precursor solution was prepared to a final concentration of 4 mg/mL (telocollagen derived from bovine skin, Cell Systems), centrifuged to remove large air bubbles and syringed into the funnel, which had been placed on an impermeable styrene sheet. A styrene rod of 1 mm diameter was oiled and mounted into a top jig piece and carefully lowered into the funnel, locking into the hole of the top membrane. After 2 h gelation at 37 °C, the funnel assembly was placed on pre-wetted paper towels and the styrene pulled out from between the funnel and paper towels to begin water absorption from the collagen gel. Densification was undertaken at room temperature overnight. Once densified, membranes were removed with a razor blade; styrene rod and densified collagen tube were removed together and placed on a rotational air-drying apparatus for ∼3 h. Once completed, the densified collagen tube was removed from the rod and transferred to a 2 mM genipin solution for 48 h, prior to washing in PBS.

### Protamine sulfate heparinisation and binding efficiency characterisation

Heparinisation of genipin-crosslinked, thick-walled densified collagen grafts was undertaken modifying a published method[57] to produce a heparin coating of 100 U per 10 mm tube length. Densified collagen tubes were immersed in 10 mg/mL protamine sulfate solution for 1h at room temperature. Tubes were then briefly rinsed in PBS prior to further crosslinking with 2 mM genipin solution, to crosslink protamine sulfate to collagen surface. A further rinse of the tubes with PBS removed excess genipin and the tubes were immersed in a 100 U/mL heparin solution at pH 4.5 (volume dependent upon tube length) and placed on an orbital shaker for 5 h at 35 °C. Tubes were then rinsed in PBS to remove unbound heparin. The heparin binding efficiency was measured using a Dimethyl-Methylene Blue (DMMB) assay. To generate a standard curve, 50 µL of heparin solution was added to 1 mL of DMMB solution. We found a 25% heparin binding efficiency and so the heparin solution volume was adjusted to achieve 10 U/mL per 10 mm tube length.

### Fabrication of non-crosslinked, thin-walled densified collagen tubes

Collagen precursor solution was prepared at 5 mg/mL concentration (with or without 0.25 mg/mL HA) and cast into a cylindrical mould (stainless steel or Perspex), comprising top wider cylinder and narrower bottom cylinder. Oil or 5% bovine serum albumin (BSA) layer was applied to the inner surface, and two membranes mounted sequentially on the base with silicone sealant. The device was placed on water impermeable styrene sheet during collagen gelation. A stainless steel metal mesh (0.5 mm pores) was placed in the top wider section which anchors the collagen from above, while the membranes at the bottom anchor the collagen from below. Water was removed axially through the membranes at one end of the funnel when placed the assembly was placed on paper towels. Due to water percolation, water is removed from the top of the gel first. This leads to the top collagen densifying down onto the top metal mesh but no further. The collagen in the lower cylindrical section then densifies radially inwards, detaching from the chamber surface. After 4 days of densification, the rod, mesh and densified collagen tube were extracted from the chamber, and the top disc-like part of collagen and mesh removed; the rod was subsequently pulled from the collagen tube under fluid. Gel concentration and wall thickness can be controlled by changing the diameter of the chamber along with starting collagen gel concentration.

### Dipyridamole coating and binding efficiency characterisation

To prevent surface clotting but avoid chemical crosslinking of the collagen tube surface, we elected to use a DIPY antiplatelet coating for the non-crosslinked densified collagen tubes, adapting a published method[58]. DIPY powder was dissolved in 50% ethanol:PBS at 4 mg/mL. Non-crosslinked densified collagen tubes were immersed in 20 mL of solution for 24 h (80 mg DIPY). We measured binding efficiency using ultraviolet-visible spectroscopy (UV-Vis) at 69% for densified collagen and 68% for densified collagen-HA tubes, leading to 22 µmol DIPY attached to inner and outer surfaces, and internal pore surfaces of the collagen tubes.

### Scanning electron microscopy

Densified collagen tubes were removed from the densification chamber, fixed with 4% paraformaldehyde for 24 h and washed with PBS. Samples were briefly dipped in deionised water to remove any buffer salts, and excess water was removed using filter paper. Samples were subsequently plunge-frozen in liquid ethane and transferred to liquid nitrogen-cooled brass inserts. Freeze-dried was performed overnight in a liquid nitrogen turbo freeze-drier (Quorum K775X). The samples were mounted on aluminium SEM stubs using conductive carbon sticky pads (Agar Scientific) and conductive silver paint, and coated with a 35 nm gold layer using a Quorum K575X sputter coater. The outer surface of the densified collagen tube was examined using a Verios 460 scanning electron microscope (FEI/ThermoFisher Scientific) operated at an accelerating voltage of 2 kV and a probe current of 13 pA. Secondary electron images were acquired using a Through-Lens detector (TLD) in full immersion mode. Manual measurements of fibril diameters were taken using ImageJ (Supplementary Fig. S2).

### Compliance and burst pressure measurements

A custom-made pressure measurement and control system was used, consisting of an instrumentation amplifier, an Arduino process controller and serial data logging software (Termite) – see previous publication for details[32]. For compliance measurements, the DIPY-coated, rat-sized collagen tubes (*n* ≥ 5), fabricated with and without HA, alongside native rat abdominal aorta (*n =* 5), were mounted on a Boston Scientific Sterling^TM^ PTA balloon dilatation catheter (6 mm balloon diameter, 20 mm balloon length and 80 cm shaft length). Specimens had a luminal diameter of ∼1.2 mm, wall thickness of ∼100 µm, and length of 30 mm. The balloon catheter was used to induce physiologically relevant intraluminal pressure (80-120 mmHg). Pressure was monitored using an industrial pressure sensor with an operating pressure of 50 psi (NPI-19H-050AV, Mouser Electronics) positioned downstream of the sample. A Keyence laser micrometer was used for real time measurement of the graft’s outer diameter at 80 and 120 mmHg. Compliance was calculated using the following formula:

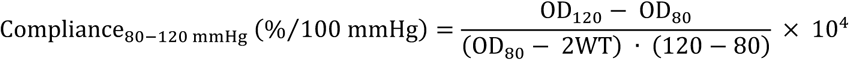

where OD120 and OD80 are the outer diameters at 120 and 80 mmHg respectively, and WT is the wall thickness.

Substantial differences in compliance variance were observed despite a common experimental protocol; therefore, the distributional properties of the data were further examined. Data in Fig. 4d are presented as raw compliance values to preserve information regarding differences in variance between groups. Compliance variance scaled approximately with the square of the mean (Supplementary Fig. S4a), consistent with multiplicative variability arising from geometric and biomechanical parameters (e.g., radius, wall thickness, elastic modulus, and pressure) and therefore with a log-normal interpretation of the data[59,60]. Consistent with this interpretation, a quantile-quantile (QQ) plot of the log-transformed compliance values showed close agreement with the theoretical normal distribution (Supplementary Fig. S4b).

Flixene ePTFE grafts (Getinge Ltd., *n* = 8), with a luminal diameter of 6 mm and a wall thickness of 1 mm, were used for comparison, despite significant differences in diameter, due to a lack of availability of ∼1 mm diameter ePTFE grafts. Grafts measuring 5 cm in length were mounted on metal hypodermic tubes and fixed in place with suture (free length of 4 cm between mount ends). The grafts were pressurised with PBS using a syringe pump (Harvard apparatus) and the pressures were recorded using an industrial pressure sensor with an operating pressure of 100 psi (NPI-19H-100AV, Mouser Electronics). Outer diameter measurements were performed as described above, and compliance values were determined using the formula described above.

For rat burst pressure measurements (*n* ≥ 5), the pressure was gradually increased until rupture, and the maximum pressure was recorded.

### Suture pullout testing

The force required for suture pullout was measured using an Instron 5544 universal testing machine (Instron, Norwood, MA, USA), equipped with a 5 N load cell, based on the principles of ISO 7198 for vascular prostheses. Tubular grafts were cut transversely into 15 mm-long segments and tested in their intact tubular configuration. Owing to the 1 mm graft diameter, testing was performed on intact tubular specimens to avoid artefacts associated with longitudinal sectioning and flattening of the graft wall. A single 8–0 polypropylene suture was passed through one wall of the DIPY-coated graft using a swaged needle, with the suture positioned 2 mm from the specimen end. Both ends of the suture were secured with masking tape and gripped in the upper fixture, while the opposite end of the tubular specimen was secured using non-slip grips. To prevent dehydration during testing, fresh PBS was applied to the samples every minute using a pipette. Crosshead displacement was measured using a linear variable differential transformer (LVDT). Tests were performed on five biological replicates per group (*n* = 5) under displacement control at a crosshead speed of 1 mm/min until failure, defined as complete suture pullout.

### In vitro hemocompatibility

To determine the blood clotting index (BCI), 4 mm circular samples were punched and placed individually into 15 mL centrifuge tubes. Coagulation was initiated by adding 100 μL of fresh rat blood, supplemented with 10% (v/v) 0.1 M calcium chloride, to the surface of the materials. After 1 hour, 3 mL of distilled water was added to the tubes to release free hemoglobin from the unclotted blood. A 200 μL aliquot of the liquid phase was subsequently transferred to a 96-well plate for UV-Vis measurement at 540 nm. The control group consisted of blood and CaCl2 mixed directly with water. The %BCI was determined using BCI (%) = AS/AC × 100 where AC denotes the absorbance of the control (100 μL of fresh rat blood mixed with 10% (v/v) 0.1 M calcium chloride solution and 3 mL of water) and AS denotes the absorbance of the sample.

To evaluate the hemolytic potential, erythrocytes were isolated from anticoagulated Sprague-Dawley rat blood (collected in Acid Citrate Dextrose). The cellular fraction was separated via centrifugation at 3000 rpm for 10 minutes, followed by two successive PBS washes to remove residual plasma. A 2% (v/v) suspension of the resulting erythrocyte sediment was prepared in PBS. Subsequently, 0.2 mL of this suspension was introduced to 10 mL of the overnight PBS-incubated sample extracts. After a final 10-minute centrifugation cycle at 3000 rpm, the supernatant absorbance was quantified at 540 nm using a UV microplate reader. The hemolysis ratio (HR) was determined by comparing sample absorbance AS against positive AP (0.1% Triton X-100) and negative AN (PBS) controls using the following formula:

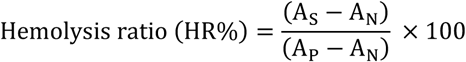

Red blood cell (RBC) attachment to the specimen surfaces was quantified through an adhesion assay. Circular samples (4 mm diameter) were inoculated with 100 µL of a 2% (v/v) erythrocyte suspension and maintained at 37 °C for 60 min. To remove non-adherent cells, 3 mL of deionized water was introduced to each vessel under mild agitation, triggering hemolysis of the remaining free RBCs. The resulting supernatant’s optical density (ODsample) was measured at 540 nm via UV-Vis spectrophotometry. A 100 µL aliquot of the 2% RBC suspension lysed directly in distilled water functioned as the control reference (ODreference). The percentage of cellular adhesion was calculated using the following formula:

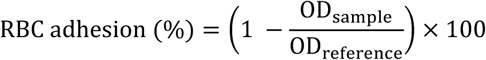

### In vivo biocompatibility study

Thick-walled genipin-crosslinked and non-crosslinked densified collagen tubes were fabricated as above, cut open longitudinally using cuticle scissors, and 5 mm diameter disc samples were obtained using a biopsy punch, excised from the lumen side outward. SMCs were cultured in a T75 flask in SMC medium (Promocell) and seeded on one side of the scaffold at 2 x 10^6^ cells/mL. Seeded samples were subsequently cultured for 1 week prior to implantation. Eighteen Sprague-Dawley rats (Charles River) were anesthetized using 4% isoflurane and maintained in 1.5% isoflurane. Samples were implanted subcutaneously in a pairwise configuration in the dorsal region. For each timepoint (1, 2, and 8 weeks), samples were implanted for both genipin-crosslinked (*n* = 6) and non-crosslinked (*n = 3)* groups. An additional group (*n* = 3) of genipin-crosslinked samples using collagen derived from rat tail, as opposed to bovine skin, was used as a control for use in case of an adverse immune reaction. At retrieval, samples were explanted together with the surrounding native tissue and fixed in 10% formalin. Following paraffin embedding and sectioning, histological analysis was performed using hematoxylin and eosin (H&E) and Masson’s trichrome staining.

### Implantation of densified collagen tubes as an abdominal aorta interposition graft

Thirteen Sprague-Dawley rats (215-340 g, Charles River) were implanted, one graft per animal (Supplementary Table 1). Rats were anesthetized using 4% isoflurane and maintained in 1.5% isoflurane. Surgeries took approximately 1.5 hours. Blood flow to abdominal aorta was interrupted with a pair of microsurgical clamps – distal to the renal arteries and proximal to the aortic bifurcation – and a 3 mm-long segment of abdominal aorta was excised, leaving 1 mm cuffs at each end. Small lumbar branches draining into the aorta were tied off using size 10’0 silk suture (BEAR Medic Corporation). Collagen grafts were measured and cut to length using surgical scissors. Suture lines were approximately 5 mm apart. Grafts were anastomosed to native aorta with 8-0 Prolene (BEAR Medic Corporation) sutures with a continuous (uninterrupted) stitch. Once both ends had been attached, the pocket was packed with hemostat (Surgicel) and the clamps removed, first distal and then proximal clamps. Surgicel was necessary to control immediate bleeding around the suture sites upon clamp removal. Surgicel was removed after 3-5 minutes to demonstrate clear pulsation of the aorta proximal and distal to the graft. Total aorta clamp time was approximately 45 min. Upon extraction of the collagen graft and peri-anastomotic tissue, the grafts were perfused with PBS and fixed using 10% formalin for 24 h. Following PBS washes, tissue samples were embedded in paraffin.

### High resolution contrast CT scan protocol

Rats were weighed and anaesthetised using isoflurane (Induction: 5% isoflurane in 2 L/min oxygen, Maintenance: 1.5-2.5% isoflurane). Iohexol (Omnipaque 300 mg I/mL). X-ray contrast agent was injected via a lateral tail vein through a butterfly cannula using a syringe pump at a rate of 0.2 mL/min. Due to rapid renal clearance of this imaging agent in rats, the CT scan as performed concurrently with the infusion. Animals were perfused with ∼0.7 mL of the contrast agent in total. The infusion was started 2 min before the scan and a CT scan (semi-circular single field-of-view, with 720 projections at 90 ms exposure per projection at X-ray power of 70 kVp, 980 µA and 1:1 binning) of approximately 1.25 min was performed during the infusion using a Mediso nanoScan PET-CT scanner. The scan was reconstructed using Filtered Back-Projection with a RamLak filter (InterView FUSION) to obtain images with a voxel size of 40 µm. Contrast agent was cleared rapidly by the rats; however, we did get clear aortic data. 4 rats were scanned (5-8 weeks post-implantation) showing patent aortic lumen in all cases (See Supplementary Fig. S1c).

### IHC Characterisation and Imaging

Paraffin-embedded tissue sections were rehydrated and underwent heat-mediated antigen retrieval using citrate buffer (pH 6.0) (Abcam). Sections were permeabilised using 0.25% Triton X-100 for 10 min, followed blocking for 1 h in 4% normal donkey serum (NDS), 1% BSA and 0.1% Triton X-100. Primary antibodies (Supplementary Table 2) were added for overnight incubation at 4 °C in blocking buffer at 1:200. Sections were washed three times using PBS with 1% BSA. Secondary antibodies (Supplementary Table 2) were added at 1:500 in blocking buffer for 2 h at room temperature, followed by washes in PBS with 1% BSA. TrueVIEW autofluorescence quenching kit (Vector Laboratories) was applied to the sections for 5 min, followed by washes in PBS for 5 min. A DAPI (6-diamidino-2-phenylindole) counterstain was applied at 1:500 dilution for 10 min. Vibrance Antifade (Vector Laboratories) and coverslip were applied to the sections which was cured overnight prior to imaging. Histological sections were imaged using a Pannoramic Confocal Digital Slide Scanner (3DHISTECH). Fluorescence IHC images were taken using a Zeiss widefield microscope (Zeiss Axio-Observer.Z1).

## AUTHOR CONTRIBUTIONS

AWJ and AEM conceived of and designed research. AWJ wrote the manuscript and prepared the figures. AEM: manuscript revision. AWJ, FC, MAAF: tube fabrication and characterisation. YL, CTG: compliance, suture retention and burst pressure measurements. AWJ, MAAF, FC, YS: Immunohistochemical staining and imaging: AWJ, MAAF, FC, YS. Surgery, implantations and live imaging: KSP, DT, AWJ, AP, NKR. Data interpretation: AWJ, AEM, MAAF, KSP, SS, MAG, ASP, MG, GJP. KSP, SS and AEM supervised the work. All authors reviewed and approved the final manuscript.

## ACKNOWLEDGEMENTS

Financial support was provided by the Engineering and Physical Sciences Research Council (EPSRC, EP/R511675/1), the Wellcome Trust Institutional Translational Partnership Award (222062/Z/20/Z) and the Medical Research Council (MRC) MR/X502844/1, MC_PC_19032). We also acknowledge support from the BHF Translational Award TA/F/24/210055. The authors are grateful to Dr Anita Petiot and Dr Kathy Boucher from Cambridge Academy of Therapeutic Sciences (CATS) for their guidance and support. A.P. was supported by an MRC award [MR/X018776/1] and S.S. by the British Heart Foundation [FS/18/46/33663]. The authors are grateful to Dr John Ong of the University of Cambridge for providing the balloon dilatation catheter used for the compliance and burst pressure testing of rat-sized grafts and vessels. We wish to acknowledge James Warner and Katie Vickers at the Histopathology Core Facility for paraffin processing and embedding, and useful discussions for characterisation. We also wish to acknowledge Martin Rice and his team at the Anne McClaren Laboratory for animal support, Louise Turvill of the Department of Pathology, University of Cambridge for histological sectioning and staining and Stefan Savage of the Department of Engineering for machining the Perspex funnels.

## ETHICS APPROVAL STATEMENT

All surgical procedures were performed in accordance with institutional guidelines and MHRC guidelines and UK Home Office Project licence (PD846B219).

## DECLARATION OF GENERATIVE AI AND AI-ASSISTED TECHNOLOGIES

ChatGPT was used to assist in generating Python plotting scripts; all outputs were reviewed and verified by the authors.

## SUPPLEMENTARY MATERIAL

## SUPPLEMENTARY MATERIAL

**Supplementary Fig. S1.**
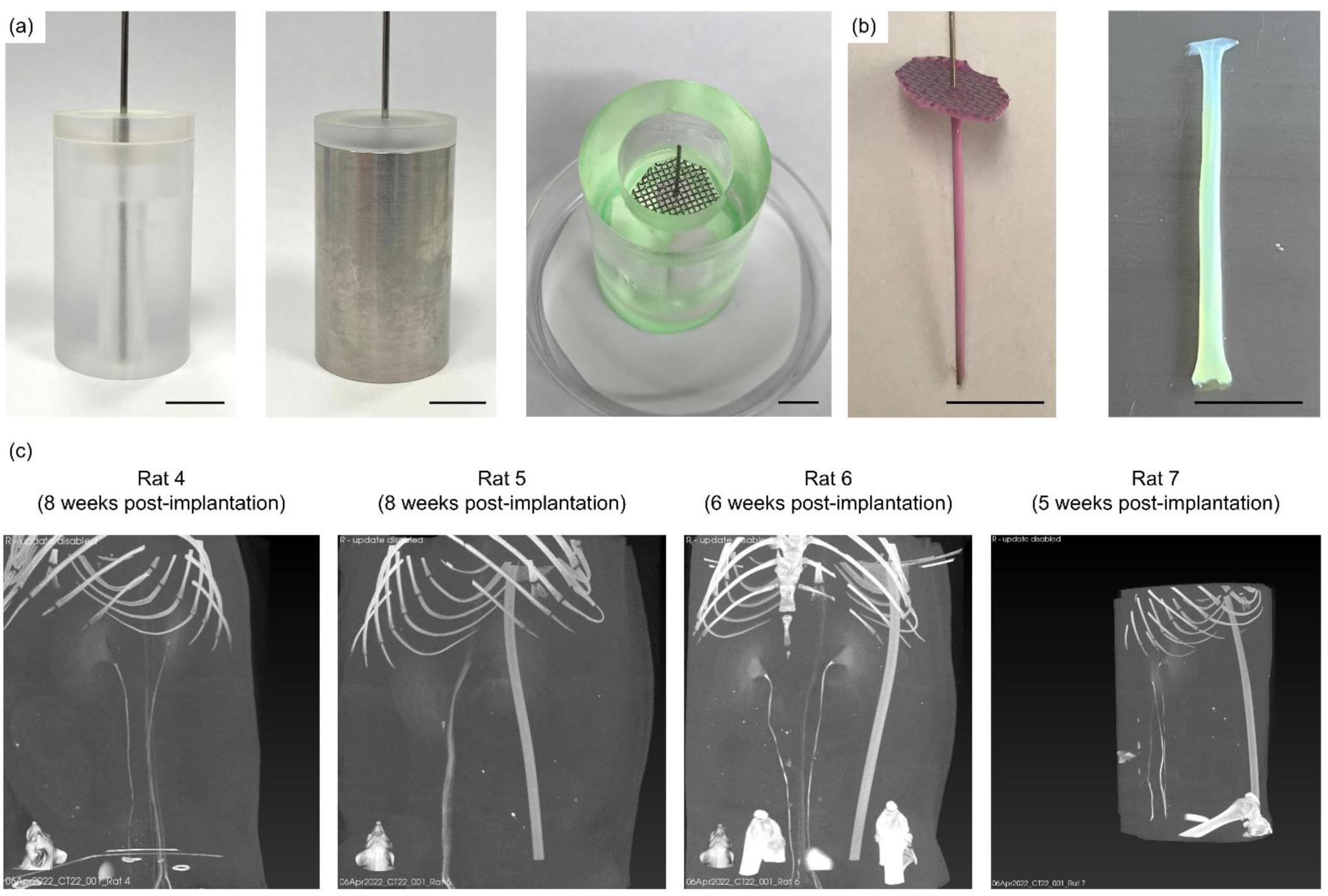
(a) Additional images of miniature chambers for constrained funnel densification. Perspex and stainless steel funnels presented (the latter for autoclaving and sterility). Also shown is the chamber with the central rod and top mesh in place, with the chamber placed on a water impermeable styrene sheet prior to gelation to prevent premature water removal through the membranes. Scale bars: 10 mm. **(b)** After densification, the mesh, rod and collagen tube are extracted from the chamber and the top collagen in the mesh is cut off. The thin-walled densified collagen tube can subsequently be removed from the rod (under fluid). Image shows densified collagen tube after treatment with DIPY for 24h. Scale bars: 10 mm. (c) High resolution contrast computed tomography scans of genipin-crosslinked tubes in vivo.

**Supplementary Figure S2.**
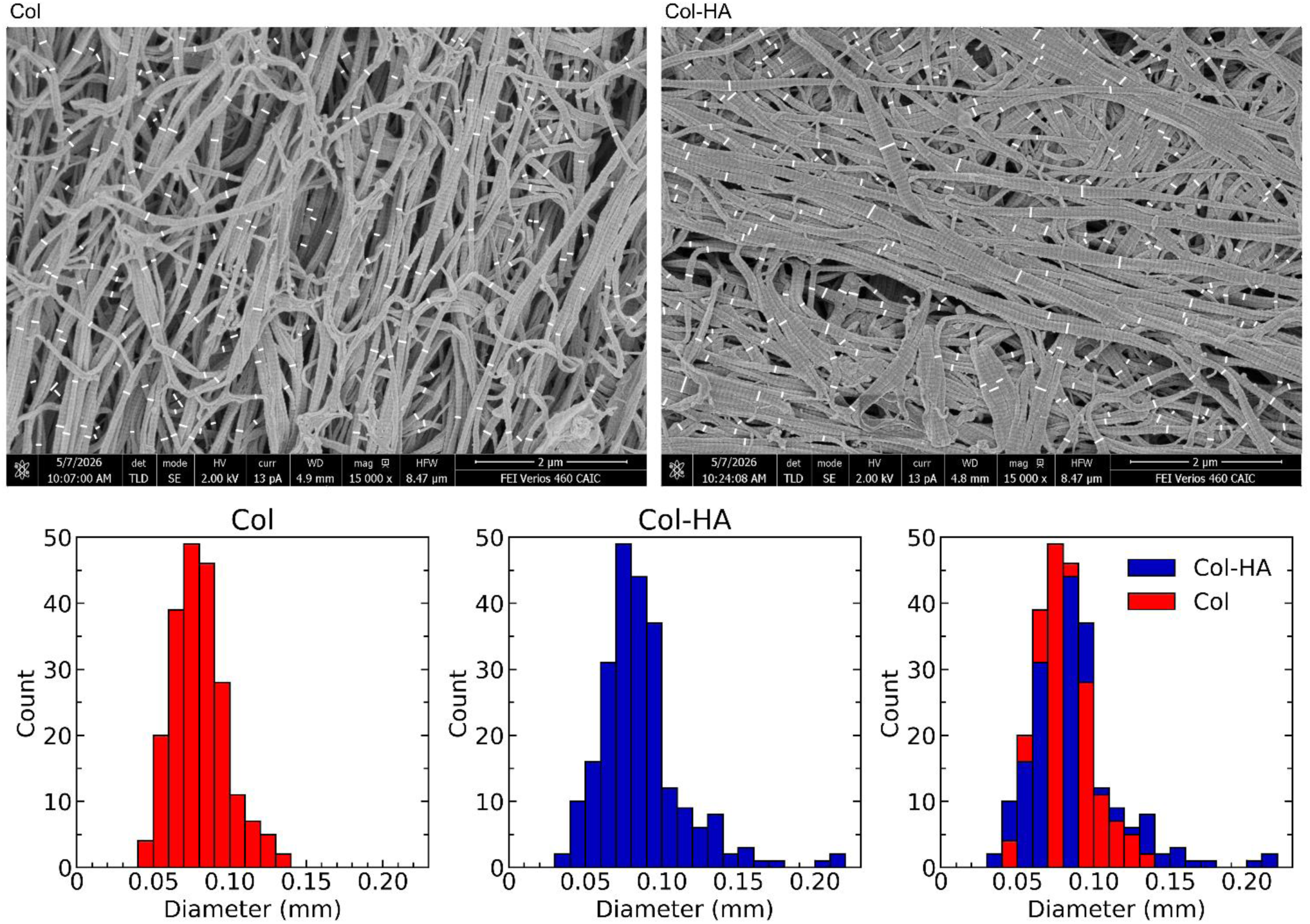
SEM fibril diameter analysis. Representative images with measurements of collagen and collagen with HA grafts. Histograms with right-closed bins of 0.010 mm.

**Supplementary Figure S3.**
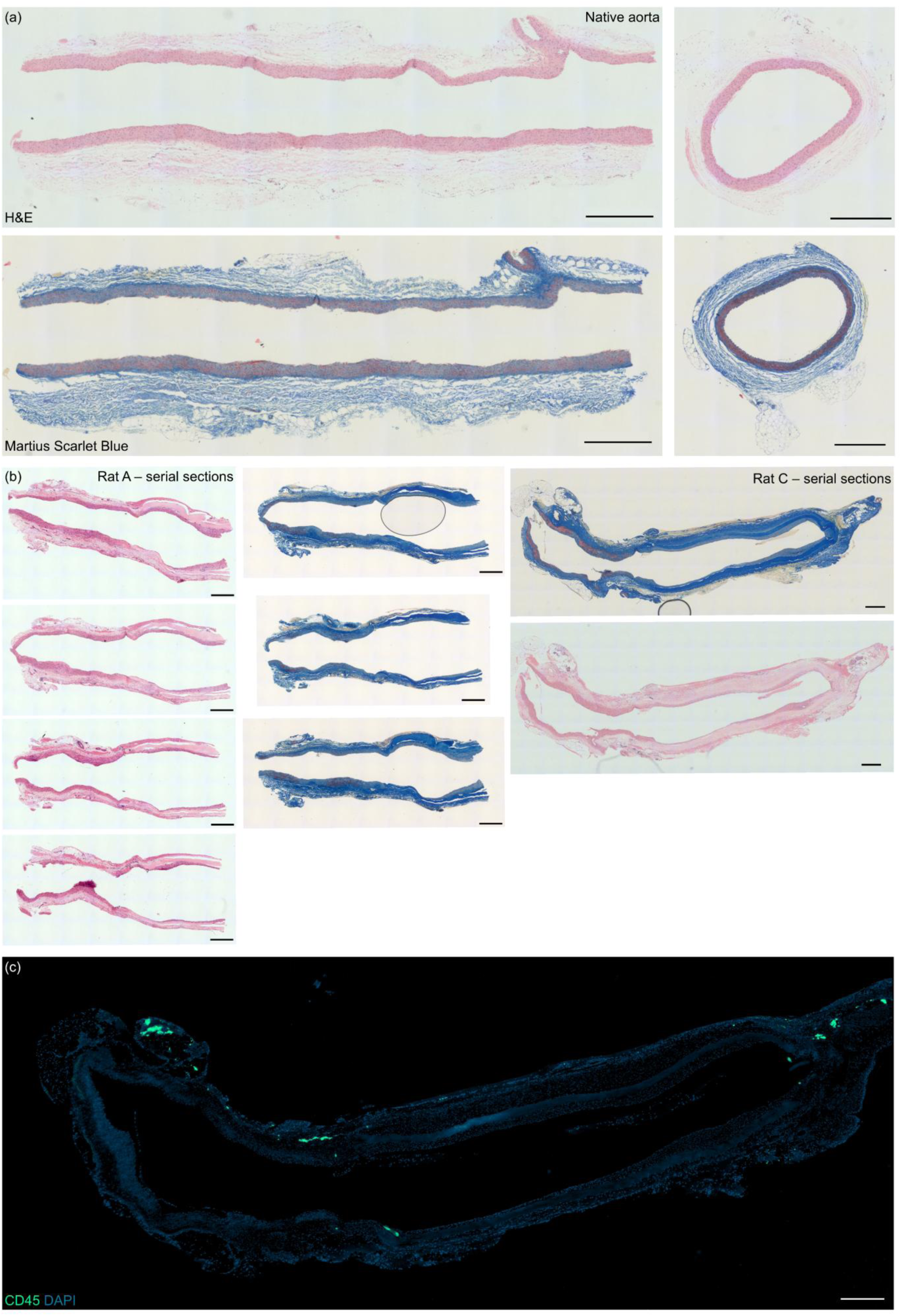
H&E and Martius Scarlet Blue histological staining of **(a)** rat abdominal aorta and **(b)** non-crosslinked collagen tubes serial sections. **(c)** Whole-section CD45 immunostaining of Rat C graft. Scalebars: 500 µm.

**Supplementary Figure S4.**
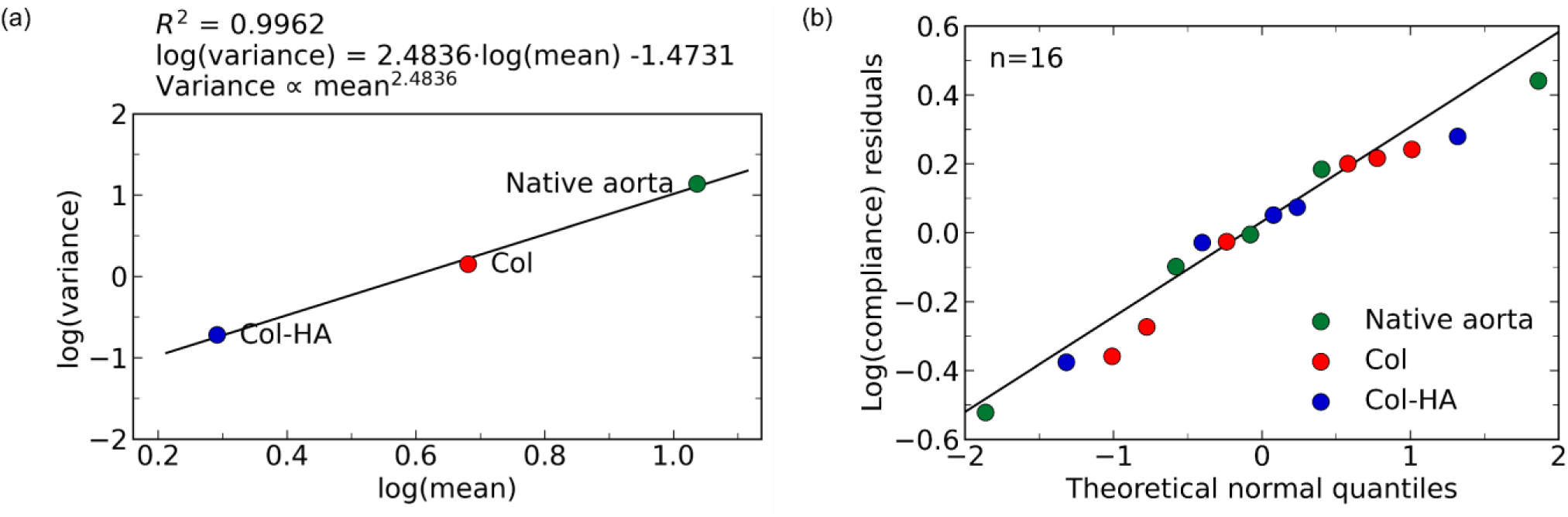
Further compliance analysis. **(a)** Linear regression of log(variance) vs log(mean) for compliance shows that variance is approximately proportional to the square of the mean, consistent with multiplicative variability and compatible with a log-normal model for compliance[59,60]. **(b)** QQ-plot of pooled log-transformed compliance residuals relative to the theoretical normal distribution. Residuals were calculated relative to the mean of each experimental group and pooled to assess whether deviations from the group means were consistent with a normal distribution.

**Supplementary Table 1.**
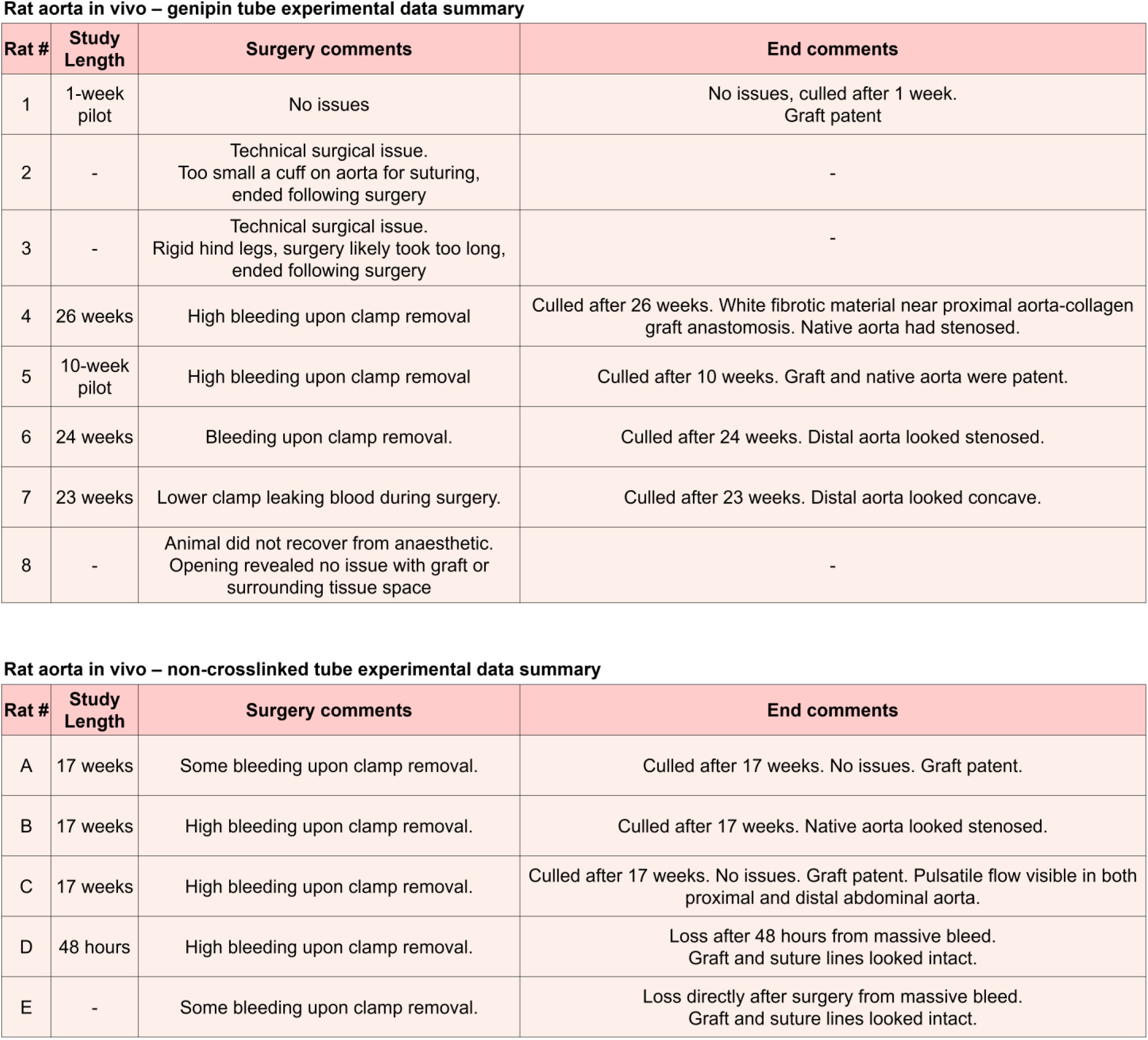
Surgical data from *in vivo* experiments.

**Rat aorta in vivo – genipin tube experimental data summary**
| Rat # | Study Length | Surgery comments | End comments |
| --- | --- | --- | --- |
| 1 | 1-week pilot | No issues | No issues, culled after 1 week. Graft patent |
| 2 | - | Technical surgical issue. Too small a cuff on aorta for suturing, ended following surgery | - |
| 3 | - | Technical surgical issue. Rigid hind legs, surgery likely took too long, ended following surgery | - |
| 4 | 26 weeks | High bleeding upon clamp removal | Culled after 26 weeks. White fibrotic material near proximal aorta-collagen graft anastomosis. Native aorta had stenosed. |
| 5 | 10-week pilot | High bleeding upon clamp removal | Culled after 10 weeks. Graft and native aorta were patent. |
| 6 | 24 weeks | Bleeding upon clamp removal. | Culled after 24 weeks. Distal aorta looked stenosed. |
| 7 | 23 weeks | Lower clamp leaking blood during surgery. | Culled after 23 weeks. Distal aorta looked concave. |
| 8 | - | Animal did not recover from anaesthetic. Opening revealed no issue with graft or surrounding tissue space | - |

**Supplementary Table 1. Surgical data from *in vivo* experiments**
| Rat # | Study Length | Surgery comments | End comments |
| --- | --- | --- | --- |
| A | 17 weeks | Some bleeding upon clamp removal. | Culled after 17 weeks. No issues. Graft patent. |
| B | 17 weeks | High bleeding upon clamp removal. | Culled after 17 weeks. Native aorta looked stenosed. |
| C | 17 weeks | High bleeding upon clamp removal. | Culled after 17 weeks. No issues. Graft patent. Pulsatile flow visible in both proximal and distal abdominal aorta. |
| D | 48 hours | High bleeding upon clamp removal. | Loss after 48 hours from massive bleed. Graft and suture lines looked intact. |
| E | - | Some bleeding upon clamp removal. | Loss directly after surgery from massive bleed. Graft and suture lines looked intact. |

**Supplementary Table 2.**
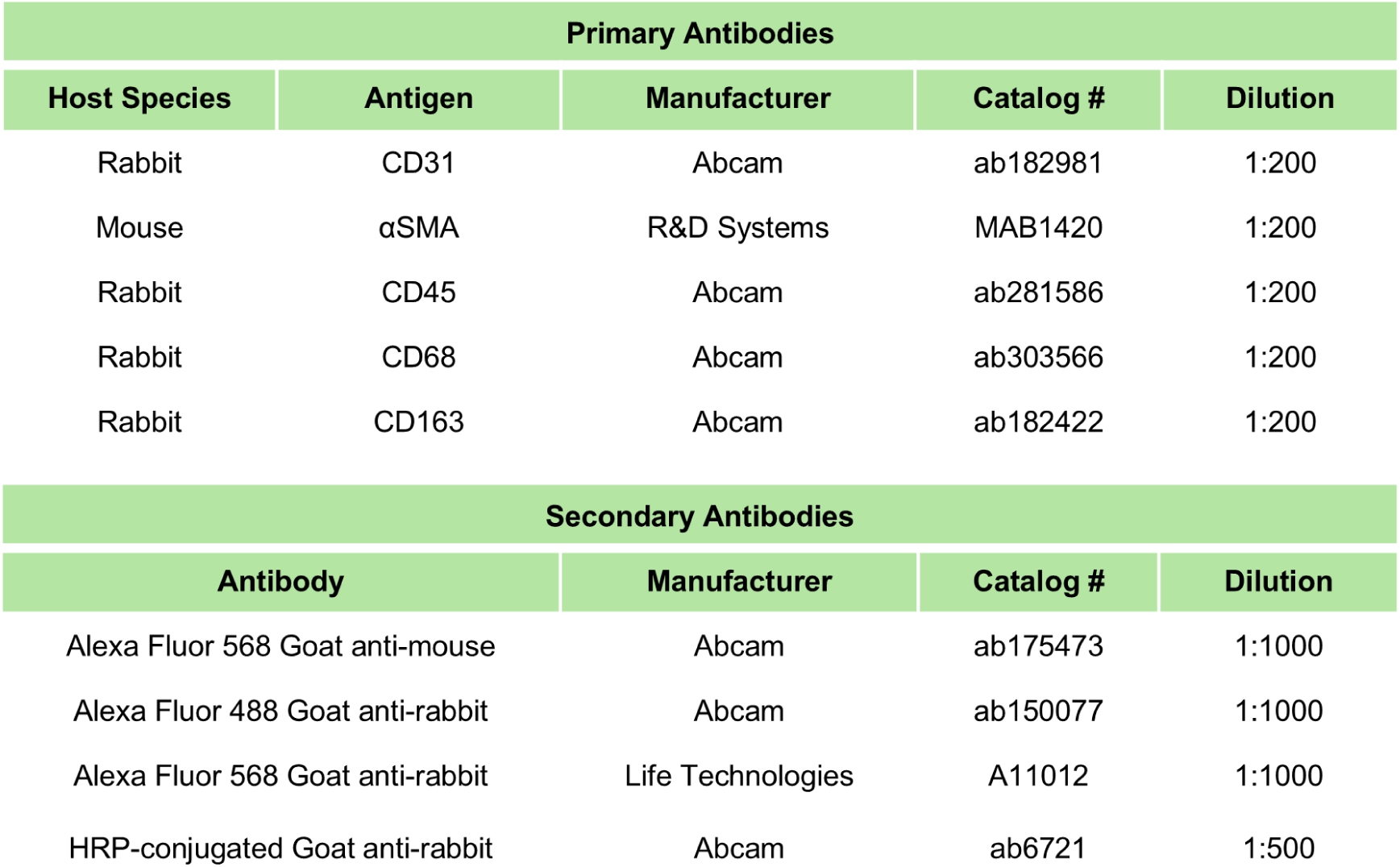
Primary and secondary antibodies.

| Primary Antibodies |  |  |  |  |
| --- | --- | --- | --- | --- |
| Host Species | Antigen | Manufacturer | Catalog # | Dilution |
| Rabbit | CD31 | Abcam | ab182981 | 1:200 |
| Mouse | αSMA | R&D Systems | MAB1420 | 1:200 |
| Rabbit | CD45 | Abcam | ab281586 | 1:200 |
| Rabbit | CD68 | Abcam | ab303566 | 1:200 |
| Rabbit | CD163 | Abcam | ab182422 | 1:200 |
| Secondary Antibodies |  |  |  |  |
| Antibody |  | Manufacturer | Catalog # | Dilution |
| Alexa Fluor 568 Goat anti-mouse |  | Abcam | ab175473 | 1:1000 |
| Alexa Fluor 488 Goat anti-rabbit |  | Abcam | ab150077 | 1:1000 |
| Alexa Fluor 568 Goat anti-rabbit |  | Life Technologies | A11012 | 1:1000 |
| HRP-conjugated Goat anti-rabbit |  | Abcam | ab6721 | 1:500 |

## REFERENCES

[1] Halbert RJ, Nicholson G, Nordyke RJ, Pilgrim A, Niklason L. Patency of ePTFE Arteriovenous Graft Placements in Hemodialysis Patients: Systematic Literature Review and Meta-Analysis. Kidney360 2020;1:1437–46. 10.34067/KID.0003502020.

[2] Leal BBJ, Wakabayashi N, Oyama K, Kamiya H, Braghirolli DI, Pranke P. Vascular Tissue Engineering: Polymers and Methodologies for Small Caliber Vascular Grafts. Front Cardiovasc Med 2021;7. 10.3389/fcvm.2020.592361.

[3] Carrabba M, Madeddu P. Current strategies for the manufacture of small size tissue engineering vascular grafts. Front Bioeng Biotechnol 2018;6. 10.3389/fbioe.2018.00041.

[4] Liu J, Song Q, Yin W, Li C, An N, Le Y, et al. Bioactive scaffolds for tissue engineering: A review of decellularized extracellular matrix applications and innovations. Exploration 2025;5. 10.1002/EXP.20230078.

[5] Golebiowska AA, Intravaia JT, Sathe VM, Kumbar SG, Nukavarapu SP. Decellularized extracellular matrix biomaterials for regenerative therapies: Advances, challenges and clinical prospects. Bioact Mater 2024;32:98–123. 10.1016/j.bioactmat.2023.09.017.

[6] Yamanaka H, Yamaoka T, Mahara A, Morimoto N, Suzuki S. Tissue-engineered submillimeter-diameter vascular grafts for free flap survival in rat model. Biomaterials 2018;179:156–63. 10.1016/j.biomaterials.2018.06.022.

[7] Wu QY, Chen CY, Li JC, Chen ZG, Chen SC, Xie C, et al. Interpenetrated hierarchical polyzwitterionic hydrogels promote decellularized small-diameter vascular grafts regeneration. Chemical Engineering Journal 2026;529. 10.1016/j.cej.2026.172764.

[8] Li Q, Ding X, Chen C, Zhang K, Dong R. An overview of small diameter vascular grafts: from materials to fabrication. Mater Adv 2025;6:6221–42. 10.1039/d5ma00663e.

[9] Wu HC, Wang TW, Kang PL, Tsuang YH, Sun JS, Lin FH. Coculture of endothelial and smooth muscle cells on a collagen membrane in the development of a small-diameter vascular graft. Biomaterials 2007;28:1385–92. 10.1016/j.biomaterials.2006.11.012.

[10] Koens MJW, Faraj KA, Wismans RG, Van Der Vliet JA, Krasznai AG, Cuijpers VMJI, et al. Controlled fabrication of triple layered and molecularly defined collagen/elastin vascular grafts resembling the native blood vessel. Acta Biomater 2010;6:4666–74. 10.1016/j.actbio.2010.06.038.

[11] Janairo RRR, Henry JJD, Lee BLP, Hashi CK, Derugin N, Lee R, et al. Heparin-modified small-diameter nanofibrous vascular grafts. IEEE Trans Nanobioscience 2012;11:22–7. 10.1109/TNB.2012.2188926.

[12] de Valence S, Tille JC, Mugnai D, Mrowczynski W, Gurny R, Möller M, et al. Long term performance of polycaprolactone vascular grafts in a rat abdominal aorta replacement model. Biomaterials 2012;33:38–47. 10.1016/j.biomaterials.2011.09.024.

[13] Gao G, Kim H, Kim BS, Kong JS, Lee JY, Park BW, et al. Tissue-engineering of vascular grafts containing endothelium and smooth-muscle using triple-coaxial cell printing. Appl Phys Rev 2019;6. 10.1063/1.5099306.

[14] Zhao L, Li X, Yang L, Sun L, Mu S, Zong H, et al. Evaluation of remodeling and regeneration of electrospun PCL/fibrin vascular grafts in vivo. Materials Science and Engineering C 2021;118. 10.1016/j.msec.2020.111441.

[15] Furdella KJ, Higuchi S, Behrangzade A, Kim K, Wagner WR, Vande Geest JP. In-vivo assessment of a tissue engineered vascular graft computationally optimized for target vessel compliance. Acta Biomater 2021;123:298–311. 10.1016/j.actbio.2020.12.058.

[16] Li W, Wu P, Zhang Y, Midgley AC, Yuan X, Wu Y, et al. Bilayered Polymeric Micro- And Nanofiber Vascular Grafts as Abdominal Aorta Replacements: Long-Term in Vivo Studies in a Rat Model. ACS Appl Bio Mater 2019;2:4493–502. 10.1021/acsabm.9b00641.

[17] Zhang L, Zhou J, Lu Q, Wei Y, Hu S. A novel small-diameter vascular graft: In vivo behavior of biodegradable three-layered tubular scaffolds. Biotechnol Bioeng 2008;99:1007–15. 10.1002/bit.21629.

[18] Vergauwen E, Tubeeckx MRL, Houben A, Van Vlierberghe S, Demolder M, De Meyer GRY, et al. Electrospun Polyurethane Vascular Grafts for Cerebral Revascularization: A Pilot Study on Rats. BioChem 2024;4:1–17. 10.3390/biochem4010001.

[19] Gao Y, Yi T, Shinoka T, Lee YU, Reneker DH, Breuer CK, et al. Pilot Mouse Study of 1 mm Inner Diameter (ID) Vascular Graft Using Electrospun Poly(ester urea) Nanofibers. Adv Healthc Mater 2016;5:2427–36. 10.1002/adhm.201600400.

[20] Soletti L, Nieponice A, Hong Y, Ye SH, Stankus JJ, Wagner WR, et al. In vivo performance of a phospholipid-coated bioerodable elastomeric graft for small-diameter vascular applications. J Biomed Mater Res A 2011;96 A:436–48. 10.1002/jbm.a.32997.

[21] Nieponice A, Soletti L, Guan J, Hong Y, Gharaibeh B, Maul TM, et al. In Vivo Assessment of a Tissue-Engineered Vascular Graft Combining a Biodegradable Elastomeric Scaffold and Muscle-Derived Stem Cells in a Rat Model. n.d.

[22] Li W, Chen J, Xu P, Zhu M, Wu Y, Wang Z, et al. Long-term evaluation of vascular grafts with circumferentially aligned microfibers in a rat abdominal aorta replacement model. J Biomed Mater Res B Appl Biomater 2018;106:2596–604. 10.1002/jbm.b.34076.

[23] Kumar VA, Caves JM, Haller CA, Dai E, Liu L, Grainger S, et al. Acellular vascular grafts generated from collagen and elastin analogs. Acta Biomater 2013;9:8067–74. 10.1016/j.actbio.2013.05.024.

[24] Li X, Xu J, Bartolák-Suki E, Jiang J, Tien J. Evaluation of 1-mm-diameter endothelialized dense collagen tubes in vascular microsurgery. J Biomed Mater Res B Appl Biomater 2020;108:2441–9. 10.1002/jbm.b.34576.

[25] Brown RA, Wiseman M, Chuo CB, Cheema U, Nazhat SN. Ultrarapid engineering of biomimetic materials and tissues: Fabrication of nano- and microstructures by plastic compression. Adv Funct Mater 2005;15:1762–70. 10.1002/adfm.200500042.

[26] Micol LA, Ananta M, Engelhardt EM, Mudera VC, Brown RA, Hubbell JA, et al. High-density collagen gel tubes as a matrix for primary human bladder smooth muscle cells. Biomaterials 2011;32:1543–8. 10.1016/j.biomaterials.2010.10.028.

[27] Li X, Xu J, Nicolescu CT, Marinelli JT, Tien J. Generation, Endothelialization, and Microsurgical Suture Anastomosis of Strong 1-mm-Diameter Collagen Tubes. Tissue Eng Part A 2017;23:335–44. 10.1089/ten.tea.2016.0339.

[28] Loy C, Lainé A, Mantovani D. Rotation-based technique for the rapid densification of tubular collagen gel scaffolds. Biotechnol J 2016;11:1673–9.

[29] Isenberg BC, Williams C, Tranquillo RT. Small-diameter artificial arteries engineered in vitro. Circ Res 2006;98:25–35. 10.1161/01.RES.0000196867.12470.84.

[30] Ban E, Franklin JM, Nam S, Smith LR, Wang H, Wells RG, et al. Mechanisms of Plastic Deformation in Collagen Networks Induced by Cellular Forces. Biophys J 2018;114:450–61. 10.1016/j.bpj.2017.11.3739.

[31] Seliktar D, Black RA, Vito RP, Nerem RM. Dynamic Mechanical Conditioning of Collagen-Gel Blood Vessel Constructs Induces Remodeling In Vitro. 2000.

[32] Justin AW, Cammarata F, Guy AA, Estevez SR, Burgess S, Davaapil H, et al. Densified collagen tubular grafts for human tissue replacement and disease modelling applications. Biomaterials Advances 2023;145. 10.1016/j.bioadv.2022.213245.

[33] Tsai C-C, Chang Y, Sung H-W, Hsu J-C, Chen C-N. Effects of heparin immobilization on the surface characteristics of a biological tissue fixed with a naturally occurring crosslinking agent (genipin): an in vitro study. vol. 22. 2001.

[34] Krzyszczyk P, Schloss R, Palmer A, Berthiaume F. The role of macrophages in acute and chronic wound healing and interventions to promote pro-wound healing phenotypes. Front Physiol 2018;9. 10.3389/fphys.2018.00419.

[35] Abbott WM, Megerman J, Hasson JE, L’Italien G, Warnock DF. Effect of compliance mismatch on vascular graft patency. J Vasc Surg 1987;5:376–82. 10.1067/mva.1987.avs0050376.

[36] Szafron JM, Heng EE, Boyd J, Humphrey JD, Marsden AL. Hemodynamics and Wall Mechanics of Vascular Graft Failure. Arterioscler Thromb Vasc Biol 2024;44:1065–85. 10.1161/ATVBAHA.123.318239.

[37] Rovas G, Bikia V, Stergiopulos N. Design and computational optimization of compliance-matching aortic grafts. Front Bioeng Biotechnol 2023;11. 10.3389/fbioe.2023.1179174.

[38] Weisel JW, Litvinov RI. Visualizing thrombosis to improve thrombus resolution. Res Pract Thromb Haemost 2021;5:38–50. 10.1002/rth2.12469.

[39] Santos EMM, Marquering HA, Den Blanken MD, Berkhemer OA, Boers AMM, Yoo AJ, et al. Thrombus Permeability Is Associated with Improved Functional Outcome and Recanalization in Patients with Ischemic Stroke. Stroke 2016;47:732–41. 10.1161/STROKEAHA.115.011187.

[40] Adolph R, Vorp DA, Steed DL, Webster MW, Kameneva M V, Watldns SC. Cellular content and permeability of intraluminal thrombus in abdominal aortic aneurysm. J Vasc Surg 1997;25:916–26.

[41] Saeidi N, Karmelek KP, Paten JA, Zareian R, DiMasi E, Ruberti JW. Molecular crowding of collagen: A pathway to produce highly-organized collagenous structures. Biomaterials 2012;33:7366–74. 10.1016/j.biomaterials.2012.06.041.

[42] Cammarata F, Al Fahad MA, Melamed S, Trajkovski D, Chadwick J, Winkel A, et al. Modulating Fibrillogenesis Enhances the Biofunctionality of Dense Collagen Hydrogel. Cell Biomaterials (Accepted) 2026.

[43] Paderi JE, Stuart K, Sturek M, Park K, Panitch A. The inhibition of platelet adhesion and activation on collagen during balloon angioplasty by collagen-binding peptidoglycans. Biomaterials 2011;32:2516–23. 10.1016/j.biomaterials.2010.12.025.

[44] Sun L, Li B, Song W, Zhang K, Fan Y, Hou H. Comprehensive assessment of Nile tilapia skin collagen sponges as hemostatic dressings. Materials Science and Engineering C 2020;109. 10.1016/j.msec.2019.110532.

[45] Siljander P, Lassila R. Studies of Adhesion-Dependent Platelet Activation Distinct Roles for Different Participating Receptors Can Be Dissociated by Proteolysis of Collagen. Arterioscler Thromb Vasc Biol 1999;19:3033–43.

[46] Zhen L, Quiroga E, Creason SA, Chen N, Sapre TR, Snyder JM, et al. Synthetic vascular graft that heals and regenerates. Biomaterials 2025;320. 10.1016/j.biomaterials.2025.123206.

[47] Cao G, Xuan X, Hu J, Zhang R, Jin H, Dong H. How vascular smooth muscle cell phenotype switching contributes to vascular disease. Cell Communication and Signaling 2022;20. 10.1186/s12964-022-00993-2.

[48] Tsai MC, Chen L, Zhou J, Tang Z, Hsu TF, Wang Y, et al. Shear stress induces synthetic-to-contractile phenotypic modulation in smooth muscle cells via peroxisome proliferator-activated receptor α/δ activations by prostacyclin released by sheared endothelial cells. Circ Res 2009;105:471–80. 10.1161/CIRCRESAHA.109.193656.

[49] Aldenhoff YBJ, Koole LH, Curtis A, Descouts P. Platelet adhesion studies on dipyridamole coated polyurethane surfaces. Eur Cell Mater 2003;5:61–7. 10.22203/eCM.v005a06.

[50] Aldenhoff YBJ, Blezer R, Lindhout T, Koole LH. Photo-immobilization of dipyridamole (Persantin®) at the surface of polyurethane biomaterials: reduction of in vitro thrombogenicity. Biomaterials 1997;18:167–72.

[51] Lichota A, Szewczyk EM, Gwozdzinski K. Factors affecting the formation and treatment of thrombosis by natural and synthetic compounds. Int J Mol Sci 2020;21:1–54. 10.3390/ijms21217975.

[52] Farndale RW, Sixma JJ, Barnes MJ, De Groot PG. The role of collagen in thrombosis and hemostasis 2004;2:561–73.

[53] Brian Savage B, Ginsberg MH, Ruggeri ZM. Influence of Fibrillar Collagen Structure on the Mechanisms of Platelet Thrombus Formation Under Flow. Blood 1999;94:2704–15.

[54] Jones B, Debski A, Hans CP, Go MR, Agarwal G. Structurally abnormal collagen fibrils in abdominal aortic aneurysm resist platelet adhesion. Journal of Thrombosis and Haemostasis 2022;20:470–7. 10.1111/jth.15576.

[55] Verrier ED, Boyle EM, Pohlman TH, Johnson MC. Endothelial Cell Injury in Cardiovascular Surgery: The Systemic Inflammatory Response. vol. 63. 1997.

[56] Kern C, Bonventre J V., Justin AW, Kashani K, Reynolds E, Siew K, et al. Necrosis as a fundamental driver of loss of resilience and biological decline: what if we could intervene? Oncogene 2025. 10.1038/s41388-025-03431-y.

[57] Noishiki Y, Miyata T. Antiadhesive Collagen Membrane with Heparin Slow Release. J Bioact Compat Polym 1987;2:325–33.

[58] Wu L, Kutas SK, Morrow BR, Hong L. Sustained release of dipyridamole from collagen membranes. J Drug Deliv Sci Technol 2023;80. 10.1016/j.jddst.2023.104180.

[59] Motulsky HJ, Head T, Clarke PBS. Analyzing lognormal data: A nonmathematical practical guide. Pharmacol Rev 2025;77. 10.1016/j.pharmr.2025.100049.

[60] Limpert E, Stahel WA, Abbt M. Log-normal distributions across the sciences: keys and clues: on the charms of statistics, and how mechanical models resembling gambling machines offer a link to a handy way to characterize log-normal distributions, which can provide deeper insight into variability and probability—normal or log-normal: that is the question. Bioscience 2001;51:341–52.

